# Resistance of mitochondrial DNA to chemical-induced germline mutations is independent of mitophagy in *C. elegans*

**DOI:** 10.1101/2022.03.28.486090

**Authors:** Tess C Leuthner, Laura Benzing, Brendan F Kohrn, Christina M Bergemann, Michael J Hipp, Kathleen A Hershberger, Danielle F Mello, Tymofii Sokolskyi, Ilaria R Merutka, Sarah A Seay, Scott R Kennedy, Joel N Meyer

**Author notes:** Kathleen A Hershberger, Biology Department, Grinnell College, Grinnell, IA, 50112, USA.

## Abstract

Mitochondrial DNA (mtDNA) is prone to mutation in aging and over evolutionary time, yet the processes that regulate the accumulation of *de novo* mtDNA mutations and modulate mtDNA heteroplasmy are not fully elucidated. Mitochondria lack certain DNA repair processes, which could contribute to polymerase error-induced mutations and increase susceptibility to chemical-induced mtDNA mutagenesis. We conducted error-corrected, ultra-sensitive Duplex Sequencing to investigate the effects of two known nuclear genome mutagens, cadmium and Aflatoxin B_1_, on germline mtDNA mutagenesis in *Caenorhabditis elegans*. After 1,750 generations, we detected 2,270 single nucleotide mutations. Heteroplasmy is pervasive in *C. elegans* and mtDNA mutagenesis is dominated by C:G → A:T mutations generally attributed to oxidative damage, yet there was no effect of either exposure on mtDNA mutation frequency, spectrum, or trinucleotide context signature. Mitophagy may play a role in eliminating mtDNA damage or deleterious mutations, and mitophagy-deficient mutants *pink-1* and *dct-1* accumulated significantly higher levels of mtDNA damage compared to wild-type *C. elegans* after exposures. However, there were only small differences in overall mutation frequency, spectrum, or trinucleotide context signature compared to wild-type after 3,050 generations, across all treatments. These findings suggest mitochondria harbor additional previously uncharacterized mechanisms that regulate mtDNA mutational processes across generations.

## INTRODUCTION

Mitochondria are vital organelles that provide energy and important signaling molecules for almost every form of eukaryotic life on earth. Phylogenetic evidence strongly suggests that mitochondria share ancestry with alpha-proteobacteria and arose by endosymbiosis. Over the course of 1.45 billion years of coevolution, most genes encoding mitochondrial proteins have translocated into the nuclear genome. Nevertheless, mitochondria still maintain a unique genome that encodes proteins essential for mitochondrial function.

mtDNA mutations are implicated in diseases affecting at least one person in 5,000 (1), in addition to potential roles in many neurological and metabolic disorders, various cancers, and aging (2–4). MtDNA mutation rates are often 100 times higher than nuclear DNA mutation rates in aging and over evolutionary time (5–7). Currently, replication errors by the sole mitochondrial DNA polymerase, Pol γ, rather than oxidative stress, are thought to be the main source of mtDNA mutations (8–10). Pol γ contains 3’-5’ -exonuclease and 5’ -deoxyribose lyase activities which allows for proofreading and base excision repair during mtDNA replication (11). Pol γ has limited translesion synthesis capabilities, potentially rendering mtDNA sensitive to damage-induced mutations or copy number changes due to polymerase stalling at bulky adducts (12). Mitochondria have efficient base excision repair mechanisms, but lack nucleotide excision repair and mismatch repair machinery, which may also contribute to higher mutation rates (13).

Chemical exposure can contribute to nuclear DNA mutagenesis, but few studies have investigated the role of genotoxicants in mtDNA mutagenesis. Evidence is limited regarding the effects of exogenous stress on mtDNA damage and somatic mutagenesis (14), and to our knowledge, the effects of chemical exposure on germline mtDNA mutagenesis have yet to be investigated. This is critical to understand because the basic biology of mitochondria renders mtDNA particularly susceptible to the harmful effects of environmental toxicants and stress (15–17).

Though mitochondria are limited in DNA repair capacity, organelle dynamics play a significant role in eliminating irreparable damage and maintaining mitochondrial homeostasis. The importance of mitochondrial fission, fusion, and mitophagy in regulating mitochondrial quality are highlighted through manifestation of disease phenotypes from mutations in nuclear encoded genes (18). For example, mutations in nuclear mitophagy genes PARK2 and PINK1 are associated with Parkinson’s disease. Mitochondrial dynamics are also important for mediating response to mitochondrial stress, whether metabolic or chemical, as we have reviewed in the context of chemical stress (19). A damaged organelle can fuse with a healthy organelle, and subsequently undergo fission to create healthy daughter mitochondria. Alternatively, damaged mitochondria may also be selectively targeted for degradation via mitophagy. We have previously demonstrated that genetic disruption of these pathways in *C. elegans* results in increased sensitivity to various stressors and inability to remove ultraviolet C radiation-induced mtDNA damage (20–23). Until this study, variation in accumulation of mtDNA damage associated with mitophagy function after chemical exposure had not been investigated.

Mitophagy is also thought to regulate the selective inheritance of mitochondrial genomes in the germline, where mtDNA is, in most metazoans, uniparentally inherited. Despite the high mtDNA copy number in oocytes, only a few mtDNA genome copies are distributed into primordial germ cells, an event that is referred to as the germline mitochondrial “bottleneck” (24). Evolutionary models predict that the bottleneck increases the probability of removal of deleterious mtDNA variants (25). Whether transmission of mutant mitochondrial genomes is stochastic, permitting genetic drift, or targeted via purifying selection, is controversial (26–29).

Previous studies in *C. elegans* have demonstrated that mitophagy can play an important role in regulating the germline transmission of mtDNA variants. For example, progeny of *C. elegans* that lack functional Parkin (*pdr-1*) accumulate higher frequencies of the 3.1kb *uaDf5* mtDNA deletion compared to wild-type (30). However, the role of mitophagy in regulating the transmission of *de novo* single nucleotide mutations through the germline remains an exciting area of investigation, particularly in the context of exposure to chemicals that cause mtDNA damage. A recent study by Haroon *et al*. suggests that mitophagy may play a role in mtDNA single nucleotide mutagenesis, as *C. elegans* deficient in *pdr-1* accumulate higher frequencies of mtDNA point mutations than wild-type, though only in a Pol γ exonuclease mutant background (31). Therefore, we hypothesized that exposure to mtDNA-damaging chemicals would result in higher rates of mtDNA mutation accumulation, and that this would be exacerbated in mitophagy-deficient *C. elegans*.

*C. elegans* is a well-established organism in which fundamental knowledge of spontaneous mtDNA mutational processes and other evolutionary insights (including genotoxin-induced nuclear mutation studies) has been achieved through classical mutation accumulation line experiments (32). Two independent mutation accumulation line (MA) experiments have been conducted in wild-type *C. elegans*, though only Konrad *et al*. conducted next-generation Illumina sequencing (7, 33). A third MA experiment was conducted in the *C. elegans* mutant, *gas-1*, which renders complex I of the ETC dysfunctional, thereby resulting in increased reactive oxygen species production and reduced ATP production (34). However, no other MA approach has investigated the effect of either chemical stressors or mitophagy on mtDNA mutagenesis in *C. elegans*.

A limitation of previous MA studies in *C. elegans* is that only a few mtDNA variants that arose to high frequencies or fixation were detected, with one very recent exception (35). However, it is now known that most mtDNA variants are not only heteroplasmic, but exist at very low frequencies that are lower than the error rate of standard next-generation Illumina sequencing assays. Therefore, we used a novel sequencing approach, Duplex Sequencing, that permits accurate detection of mtDNA variants at frequencies as low as one mutation in ∼10^8^ base pairs (36). Using this ultra-sensitive, mtDNA-targeted sequencing approach, we were able to detect more heteroplasmic mtDNA mutations at lower frequencies than ever before reported in *C. elegans*, and also investigate the mutational rates, spectrum, context, and genomic sites of mutagenesis after 50 generations of mutation accumulation.

To investigate the effects of chemical stress and the role of mitophagy in the origin and transmission of mtDNA mutagenesis in *C. elegans*, we performed mutation accumulation experiments in wild-type and two mitophagy mutant strains, *pink-1* and *dct-1*, under various environmental conditions. All three strains were bottlenecked for 50 generations in control conditions, in addition to chronic exposure for 50 generations to the heavy metal pollutant, cadmium (Cd, in the form of cadmium chloride, CdCl_2_) or the mycotoxin aflatoxin B_1_ (AfB_1_), both known nuclear mutagens and human carcinogens. Cd inhibits many DNA repair enzymes and interferes with antioxidant enzymes, resulting in increased levels of ROS that can damage mtDNA (37, 38). AfB_1_ metabolites form bulky DNA adducts, which can inhibit replication and result in somatic mutations in the nuclear genome (39). The accumulation of AfB_1_ in mitochondria, in addition to lack of nucleotide excision repair (NER), results in higher levels and greater persistence of mtDNA lesions compared to nuclear DNA damage, as we and others have previously measured (40–42). Therefore, we used these two toxicants as models to investigate how two different mechanisms of mtDNA damage may affect mtDNA mutational processes.

Duplex Sequencing of *C. elegans* mtDNA revealed a strong signature of oxidative damage. Exposure to levels of CdCl_2_ and AfB_1_ that caused mtDNA damage did not increase the frequency of mtDNA single nucleotide mutations (SNMs) in wild-type *C. elegans*. Surprisingly, inhibiting mitophagy did not result in an increase in mtDNA mutations in control conditions or after exposure to CdCl_2_ or AfB_1_. These results suggest that the mitochondrial genome harbors robust mechanisms of avoiding chemical-induced mtDNA mutagenesis that are independent of mitophagy.

## MATERIAL AND METHODS

### *C. elegans* strains and maintenance

This work used the *C. elegans* wild-type Bristol N2 strain and two mitophagy deficient strains, *dct-1* and *pink-1*. The *dct-1*(tm376) mutant harbors a 912 bp deletion in the promoter region of *dct-1* (DAF-16/FOXOControlled, germline Tumour affecting-1, putative orthologue to mammalian mitophagy receptor, BNIP3) and has been characterized previously (43). The *pink-1*(1779) mutant harbors a 350 bp deletion in *pink-1* (PTEN-induced kinase 1) and exhibits altered mitochondrial morphology (Luz 2015). These deletion strains were acquired from the National Bioresource Project (Tokyo, Japan), genotyped, and backcrossed into N2 six times (*dct-1*) and eight times (*pink-1*) prior to any experiments. All *C. elegans* were maintained following standard procedures (44). We replaced the potassium phosphate in traditional nematode growth medium (NMG plates) with KCl (“K-agar plates”) in order to prevent buffering which reduces the bioavailability of CdCl_2_ (45, 46). All strains were maintained at 20°C on OP50 *E. coli*.

### CdCl_2_ and AfB_1_ treatment

Stocks of CdCl_2_ and AfB_1_ (Sigma-Aldrich, St. Louis, MO) were made in ddH_2_O and DMSO vehicles, respectively, and stored at 4°C. Treatment plates were always made fresh: OP50 was spiked with either CdCl_2_ and AfB_1_ immediately prior to seeding, and plates were prepared two days prior to experiments. 100µL of OP50 was seeded on 6cm plates (containing 8mL of K-Agar) and 300µL of OP50 on 10cm plates (containing 20mL K-agar).

### Mitochondrial morphology

To assess effects of exposure on mitochondrial morphology, wild-type *C. elegans* harboring an extrachromosomal array P*myo-3*::GFP; which expresses GFP in the mitochondrial matrix of body wall muscle cells, were exposed to 10µM and 50µM of CdCl_2_ and 2µM and 10µM AfB_1_. Images were taken on a Keyence BZ-X700, and analyzed in ImageJ using the Blinder software (47). Blinded images were analyzed qualitatively by the established classification system as previously described (48), with Class 1 indicating highly networked, fused mitochondrial morphology, and Class 5 indicating extremely fragmented mitochondrial morphology. Two experimental replicates were conducted, and the total number of individuals analyzed is displayed above each stacked bar plot in Supplementary Figure S5. Statistical differences in the distribution of classes of mitochondrial morphology were determined by Fisher’s Exact Test followed by Bonferroni correction for multiple comparisons.

### Mitochondrial respiration

Age-synchronized L1 *C. elegans* were transferred to plates seeded with OP50 containing 50µM and 100µM CdCl_2_, or 10µM and 25µM AfB_1_. Approximately 48 hours later, L4 *C. elegans* were washed off the plates and respiration parameters were quantified using the Seahorse Extracellular Flux Bioanalyzer, as previously described (49). The number of individual nematodes per well were counted, such that the Oxygen Consumption Rate (OCR) measurements were normalized per worm. The mean of technical replicates per plate was then determined. Two to six independent experiments were conducted per strain per treatment. Two-way ANOVAs were run for each chemical in order to compare Cd to control and AfB_1_to control across three strains.

### Life history trait analyses

Growth was measured in three independent experiments in the ancestral (G0) wild-type, *dct-1*, and *pink-1* strains on control and treatment plates. About 300 age-synchronized L1s were plated on each 10cm plate with the following conditions: control, 50, 200, 1000µM CdCl_2_, and 10, 50, and 200µM AfB_1_. After 48 hours, L4 *C. elegans* were washed off the treatment plates and transferred to unseeded plates. Images of each plate were then captured with a Keyence BZ-X700 using brightfield, and analyzed in ImageJ with the Fiji plug-in WormSizer (51). Length data were normalized to the mean control length of each strain for each biological replicate. Box and whisker plots of length distributions were plotted in R and the means of biological replicates were used for statistical analysis (2-way ANOVA, Tukey’s HSD post-hoc analysis). Total brood size was also determined in all three strains on control, 50µM CdCl_2_ and 10µM AfB_1_ plates (N = 9-10 individuals per strain per treatment). On day 1 of adulthood, each individual worm was transferred to a new 6cm plate with the respective treatment until reproduction ceased. Plates containing offspring were stored at 20°C for two days, and then counted.

Population growth rate was measured on all MA lines at G0 and G50 to measure the effects of mutation accumulation on fitness. Population growth rate was determined by an eating race experiment as originally described by Hodgkin and Barnes (52). Each plate was monitored hourly up to the time when all bacteria had been consumed and the population dispersed. As an additional fitness measurement, total reproduction and lifespan were also determined on a subset of 20 MA lines per strain per treatment at G50. The 20 MA lines were determined with a random number generator. Of these 20 sublines, three L4 individuals (G50 sisters) were picked onto individual control plates. Founders were transferred to a new plate daily until cessation of egg-laying, and total brood was counted as described above. Biological replicates were then averaged per MA line. Individuals that were sterile were included in our analysis. Lifespan was determined by scoring individuals every day until they stopped responding to touch. Reproduction and lifespan results were censored if the animal was lost, dehydrated, or bagged. This only occurred in four of the 180 MA lines.

### mtDNA copy number and damage

OP50 was spiked with either CdCl_2_ or AfB_1_, such that the final concentrations of chemicals in the bacterial lawn were 50µM of CdCl_2_ and 10µM of AfB_1_. Plates were seeded with 100µL of each treatment, and control, and allowed to dry for two days. 20-30 gravid adults of each strain, wild-type, *pink-1*, and *dct-1*, were then picked onto each treatment plate. Adults were left on each plate for a 2-hour egg-lay, after which all of the adults were removed from the plate. After 56 hours, synchronized L4 *C. elegans* were then analyzed for mtDNA CN and damage, as described (53). Two to three biological replicates of six individual L4 *C. elegans* per strain per treatment were picked into 90µL of lysis buffer, placed at -80ºC for at least overnight, then lysed at 65ºC for one hour, followed by 95ºC for 15 minutes to inactivate Proteinase K. Long-amplicon PCR was then performed (53), with one modification that the final reaction volume was halved (25µL instead of 50µL). mtDNA CN was measured using a plasmid-based standard curve and real-time PCR as described (54). Long-amplicon PCR products were normalized to mtDNA copy number and mtDNA lesions were calculated relative to the control within each strain. Each biological sample was run in duplicate or triplicate. This experiment was performed twice. A two-way ANOVA followed by Tukey’s HSD post-hoc analysis was used to determine statistical differences in mtDNA lesions between strains and treatment. Results of mtDNA lesions following exposure to additional concentrations of CdCl_2_ and AfB_1_ that were not used in the mutation accumulation line experiment are included in Supplementary Materials (Supplementary Figure S1). The effects of exposure on mtDNA CN are described in Supplementary Figure S2.

### Mutation Accumulation Lines

Our criteria to determine a single concentration of CdCl_2_ and AfB_1_ to use for mutagenesis experiments were: 1) the concentration caused significant mtDNA damage, but 2) did not have significant effects on organismal fitness (growth, fecundity, and mitochondrial toxicity). Once we determined a concentration for each exposure, we performed a mutation accumulation line experiment in wild-type, *pink-1*, and *dct-1 C. elegans*, in control conditions, 50µM CdCl_2_, or 10µM AfB_1_ (Figure 2). One random young adult individual of each strain was isolated (G0) and allowed to reproduce. Once the offspring reached gravid adulthood, about 20 individuals were picked onto a plate. After a 2-hour egg lay, the adults were picked off the plate. When these offspring reached L4, one individual (G1) was randomly picked to an individual plate to begin the mutation accumulation experiment. The MA experiment began with 50 individual lines per strain per treatment. All lines were maintained at 20ºC. Every four days, one individual L4 was randomly picked and transferred to a new plate to propagate the next generation. This population bottlenecking was conducted for 50 generations per strain per treatment, resulting in a total of 2,500 generations of mutation accumulation per strain per treatment. The previous generation was always maintained at 15ºC, such that if an individual was sterile or dead, a back-up “sister” individual could be transferred to a new plate to continue the MA line. As in previous *C. elegans* MA experiments, if three attempts were not viable, the line was then considered extinct (55). At the end of the experiment, the average percentage of extinctions per strain per treatment was 4.5% (Supplementary Figure S3). All 450 MA lines were cryopreserved every 5 generations for 50 generations.

### DNA isolation, Duplex Sequencing library preparation, and sequencing

After 50 generations of mutation accumulation, an individual L4 from each MA line was transferred onto a control 6cm plate. As has been previously described in other MA experiments, as soon as all of the OP50 had been consumed and the population was composed largely of synchronized L1s, the *C. elegans* were immediately washed off of the plate with ddH_2_O, transferred to a 1.7mL Eppendorf tube, pelleted, and flash frozen in liquid nitrogen (33–35). Total DNA was isolated with the DNeasy Blood & Tissue kit (Qiagen, Germany) following the manufacturer’s instructions. DNA quantity and purity was analyzed using a Thermo Scientific™ NanoDrop™ One Microvolume UV-Vis Spectrophotometer (Thermo Fisher Scientific) and Qubit 2.0 (Perkin Elmer Victor x2).

Illumina Sequencing libraries were prepared following the original Duplex Sequencing (DS) protocol with slight modifications (56, 57). An input of 50 ng of total DNA was used for library preparation. Total DNA was sheared (Covaris E210), followed by preparation and repair of DNA fragments using the NEBNext Ultra II Library prep kit, in which 2µL of 15µM of the unique DS adapters were added directly to the Ultra II Ligation Master Mix, according to manufacturer’s instructions (New England Biolabs). Relative mtDNA copy number of the post-adapter ligated sample was then determined via RT-qPCR (Applied biosystems StepOnePlus). This critical step determined the amount of post-ligated DNA to use as input for the pre-capture indexing PCR to optimize the number of Duplex Consensus Sequences formed for each Unique Molecular Identifier (57). The optimal family size was determined by earlier sequencing of a *C. elegans* wild-type sample compared to a known reference standard in order to determine volume of post-adapter-ligated for pre-capture indexing PCR. On average, 0.338µL of template was required (range of 0.164 – 0.632). After the pre-capture indexing-PCR, the entire product was lyophilized with the addition of uniquely designed blocking oligos. An enrichment of the *C. elegans* mitochondrial genome was then performed following the IDT xGen Hybridization Capture of DNA libraries for NGS target enrichment protocol with a custom designed Discovery Pool probe panel that covered 13,091bp of the 13,991bp *C. elegans* mitochondrial genome, capturing 93.5% of the genome. The AT-rich, highly repetitive non-coding region of the genome was excluded in order to minimize sequencing fragments that would not accurately map to the reference genome. Post-capture PCR and clean-up were then performed using according to manufacturer’s instructions. Libraries were sequenced on an Illumina NovaSeq 6000 platform to obtain 150bp paired-end reads (20 million per library). The DS adapter sequences, qPCR and Illumina primer sequences, blocking oligo sequences, and the *C. elegans* mitochondrial genome custom probe panel oligonucleotide sequences are provided in the Supplementary Data File.

### Bioinformatics processing and analysis

Sequencing data were processed on the Duke Computer Cluster using the custom Duplex-Seq Pipeline (v1.1.4) workflow developed by the Kennedy Lab (github.com/Kennedy-Lab-UW/Duplex-Seq-Pipeline) and described in detail in Supplementary Figure 3 of Kennedy et al. (56) and again in Sanchez-Contreras et al. (10). Only reads mapping to the *C. elegans* mitochondrial reference genome were analyzed, and only single nucleotide mutations were analyzed, while insertions and deletions were ignored. Possible artifacts were removed by 12-bp end clipping at the beginning of each read. A minimum of three reads were required to call a variant, with a minimum heteroplasmy cutoff of 70% per Duplex Consensus Sequence (DCS) compared to the reference genome. Other input parameters are included in the Configuration File (Supplementary Data File). A summary output file of mutations, sequencing depth, and genome coverage per library is included (Supplementary Data File). Sequencing data is uploaded and can be accessed at SRA SRP350474 (PRJNA787252). We detected three polymorphisms that were completely fixed in our *C. elegans* strains compared to the reference genome (NC_001328.1), two of which have been previously detected (35). These polymorphisms were not included in our analysis and are listed in Supplementary Table S1.

### Mutation frequency and spectrum analysis

Each mutation was only called once at each genomic position. Overall mutation frequency was calculated for each library with the following equation: [total number of unique mutations] / [total number of error-corrected nucleotides sequenced]. For mutational signature calculations, the total number of each type of the six possible transition and transversion mutations was divided by the coverage of the reference nucleotides sequenced (i.e.: [total # C:G → A:T mutations] / [total number of cytosines + guanines sequenced]). Mean mutation frequencies were calculated for each strain and treatment, followed by parametric statistical analysis and appropriate multiple comparisons corrections, as described in Results.

### Trinucleotide context mutational signature

Trinucleotide context mutations were calculated using the Bioconductor Package *MutationalPatterns* (v1.1.0) (58) after forging a *C. elegans* mitochondrial reference genome (*BSgenome* v1.58.0). The number of mutations in each of the 96 possible trinucleotide contexts were determined in reference to pyrimidines C and T, hence six possible mutation types instead of 12. The number of each trinucleotide context mutation was normalized to the average number of wild-type control mutations in each context. A Welch Two Sample T-test was performed on a specific mutation only after a significant ANOVA was determined.

### Synonymous and nonsynonymous mutation analysis

The dNdS ratio was calculated in order to determine if mtDNA evolution departed from neutrality in our MA experimental design. We used the *R* package *dNdScv*, a statistical modeling approach that allows for normalization to depth of coverage as a covariate, and incorporation of the invertebrate mitochondrial genetic code (translation table 5) (59).

### Statistical analysis

All statistical analysis and data visualization were conducted in RStudio (v1.1.463; *R* v4.0.3). Values are conveyed as means and standard error, unless otherwise indicated. We performed Welch Two Sample T-test to determine significance between two groups, and ANOVA (one way if more than two groups and two way if multiple factors, as indicated in further detail in the figure legends) followed by Tukey’s HSD to correct for multiple comparisons. Fisher’s Exact Test was performed to determine variation in the proportion of each type of mutation (Supplementary Figure S4).

## RESULTS

### CdCl_2_ and AfB_1_ exposure causes mtDNA damage in wild-type *C. elegans*

We have previously shown that exposure to CdCl_2_ and AfB_1_ results in significantly higher levels of mtDNA damage than nuclear DNA damage in wild-type *C. elegans* (40). In order to determine a single dose of CdCl_2_ and AfB_1_ to use for mutation accumulation lines and sequencing where there was detectable mtDNA damage but no evident effect on fitness, we first conducted a dose response and quantified mtDNA damage (Supplementary Figure S1), followed by measuring growth, reproduction, mitochondrial morphology, and mitochondrial respiration. Exposure to 50µM CdCl_2_ induced 0.92 lesions/10kb, and 10µM AfB_1_ induced 0.25 lesions/10kb relative to control in wild-type *C. elegans* (*P* < 0.01, Figure 1A). These concentrations of CdCl_2_ and AfB_1_ did not result in statistically significant growth delay (8.3% and 6.3% growth inhibition, respectively, (Figure 1B) or fecundity (Figure 1C)). We also observed little effect on mitochondrial morphology (Supplementary Figure S5) and no decrease in mitochondrial function (Supplementary Figure S6). Therefore, these concentrations were used for the mutation accumulation experiments in order to avoid any effects of fitness on mutation rate calculations.

**Figure 1.**
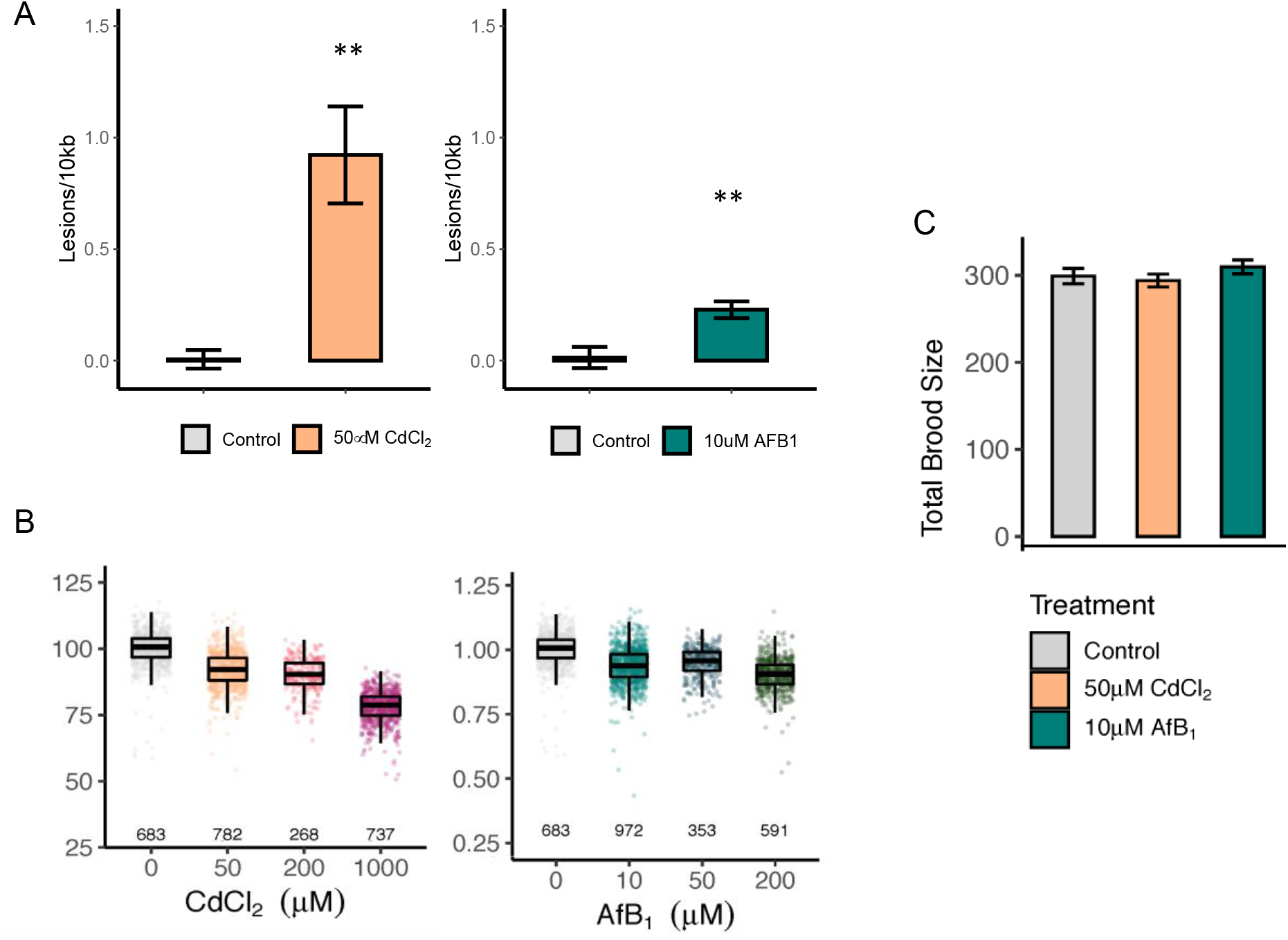
50µM CdCl_2_ and 10µM AfB_1_ exposure induces mtDNA lesions in wild-type *C. elegans*, but has no effect on growth or reproduction. (A) mtDNA damage from pools of six individual age-synchronized L4 *C. elegans* was quantified after chronic exposure to control (N = 8), 50µM CdCl_2_ (N = 8), or 10µM AfB_1_ (N = 4). (** *P* < 0.01, one-way ANOVA) (B) A dose-response was conducted to measure effects on growth after 48 hours of exposure (L1-L4). Each individual worm length was first normalized to the mean control length for each experiment. Dots represent technical replicates (individual nematodes, total N displayed) and the boxplots display the median and upper and lower quartiles. The mean normalized length was then calculated within each experimental replicate. There was no effect of growth after exposure to 50µM or 200µM CdCl_2_, but we observed 22% growth inhibition at 1,000µM CdCl_2_ compared to control (*P* = 0.15, *P* = 0.2, *P* = 0.009, respectively; one-way ANOVA, Tukey HSD). We did not observe growth inhibition after 10µM or 50µM AfB_1_ compared to control, but did observe 11% growth inhibition at 200µM AfB_1_, though trending (*P* = 0.27, *P* = 0.59, *P* = 0.09, respectively; one-way ANOVA, Tukey HSD). (C) There is no effect of either 50µM CdCl_2_ or 10µM AfB_1_ on total brood size compared to control (N = 9-10, *P* = 0.9, *P* = 0.63, respectively; one-way ANOVA, Tukey HSD). Error bars indicate standard error (* *P* < 0.05; ** *P* < 0.01).

**Figure 2.**
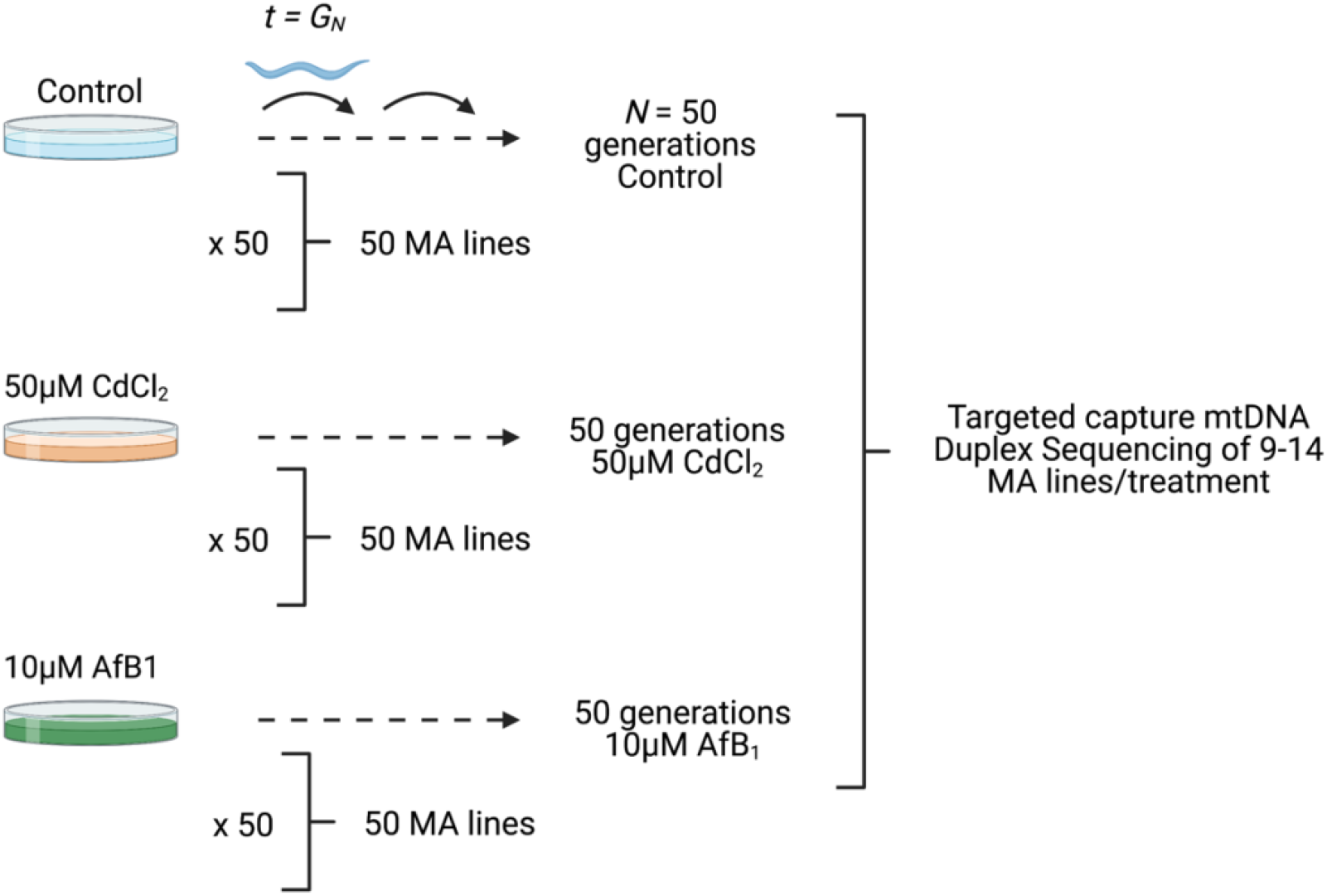
Schematic of Mutation Accumulation line experimental design. Offspring of a single founding ancestor (G0) were isolated onto individual plates. Every t = 4 days, a single L4 nematode was randomly selected and transferred to a new plate. This was repeated every generation (G) for 50 generations. We conducted MA lines on control plates, and plates that contained OP50 that was spiked with a final concentration of 50µM CdCl_2_ and 10µM AfB_1_. We conducted MA line experiments in wild-type *C. elegans* and two mitophagy mutant strains, *dct-1* and *pink-1*. 50 replicates were passaged for each strain*treatment. MA lines were randomly selected after 50 generations (9-14 MA lines/strain/treatment) for life history analysis and targeted mtDNA Duplex-Sequencing. Image created with BioRender.com.

### Wild-type *C. elegans* exhibit resistance to mtDNA damage-induced single nucleotide mutations

The strong increase in mtDNA damage suggested that mutations should be correspondingly increased. To test this hypothesis, we conducted Duplex Sequencing after 50 generations of mutation accumulation to determine the frequency and spectrum of mtDNA single nucleotide mutations on wild-type *C. elegans* in control and genotoxicant-treated conditions. To verify the Duplex Sequencing method in *C. elegans* (which had not previously been conducted before, prior to the recent work by Waneka et al. (35)) we first optimized the protocol in wild-type *C. elegans* after exposure to 7 J/m^2^ ultraviolet C radiation, where we observed a 1.6-fold increase in mtDNA SNMs compared to a control sample (4.89 versus 3.07 × 10^−5^ mutations per base pair) (Supplemental Figure S7). This confirmed that Duplex Sequencing is not only feasible in *C. elegans*, but has the sensitivity to detect effects of mtDNA damage on mutagenesis.

After confirming the utility of Duplex Sequencing to detect rare variants and exposure differences in *C. elegans*, we sequenced 11 wild-type MA lines and identified a total of 760 SNMs (an average of 70 mutations/MA line) under control conditions. The overall mtDNA mutation frequency of wild-type *C. elegans* was 10.1 × 10^−7^ SNMs/bp. Despite significant mtDNA lesions (Figure 1A), there was no effect of CdCl_2_ (14 MA lines, 874 total SNMs) or AfB_1_ (10 MA lines, 636 total SNMs) on the overall mtDNA SNM frequency in wild-type *C. elegans* (9.75 × 10^−7^ and 9.47 × 10^−7^, respectively) (Figure 3A, Supplementary Table S2). The mutation spectrum of *C. elegans* was dominated by C:G → A:T and C:G → G:C transition mutations, and C:G → T:A mutations. There was no effect of CdCl_2_ or AfB_1_ - induced mtDNA damage on mutation spectrum (Figure 3B; Welch Two Sample T-test). Wild-type *C. elegans* exhibited no strand asymmetry in mtDNA mutation accumulation in control conditions (Figure 3C, top panel). There was no strand asymmetry after exposure to CdCl_2_ (Figure 3C, middle panel). We did observe a trend of a slightly higher frequency of C → T over G → A transversions after exposure to AfB_1_ compared to control conditions (*P* = 0.08; two-way ANOVA) (Figure 3C, bottom panel).

**Figure 3.**
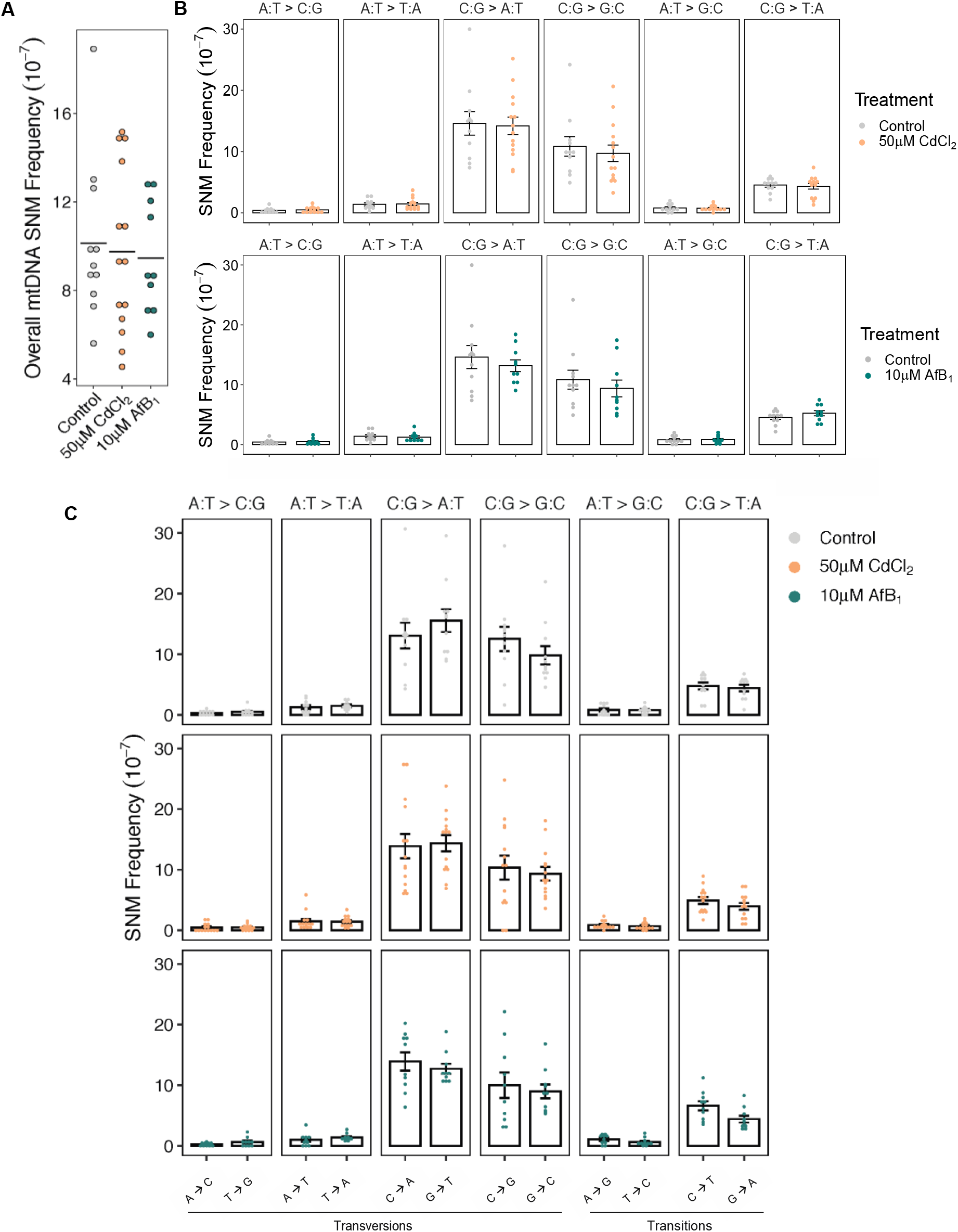
The mtDNA single nucleotide mutation signature in wild-type *C. elegans* is consistent with oxidative damage and demonstrates resistance to point mutations caused by CdCl_2_ and AfB_1_ mtDNA lesions. (A) Overall SNM frequency was determined by Duplex Sequencing after 50 generations of mutation accumulation in wild-type *C. elegans* in control conditions (gray dots; N = 11), as well as after 50 generations of exposure to 50µM CdCl_2_ (gold dots; N = 14) and 10µM AfB_1_ (green dots, N = 10). There was no effect of either 50µM CdCl_2_ or 10µM AfB_1_ on overall mtDNA mutation frequency compared to control (*P* = 0.96, *P =* 0.90, respectively); one-way ANOVA, Tukey HSD). Horizontal lines indicate mean values. (B) Mutation spectrum of control, CdCl_2_, and AfB_1_ -treated MA lines. Each dot represents a single MA line. The wild-type *C. elegans* mtDNA mutational signature was dominated by C:G → A:T and C:G → G:C transversion mutations, and there was no effect of Cd or AfB_1_ exposure on mtDNA mutational signature (Welch Two Sample T-test). Error bars represent standard error of the mean. (C) mtDNA lesions may accumulate disproportionately on mtDNA strands, resulting in mtDNA mutation strand bias. We observed a trend towards an increase in C → T over G → A mutations after exposure to AfB_1_ compared to control (*P* = 0.08; two-way ANOVA, Tukey HSD).

### Mitophagy deficiency exacerbates mtDNA damage but not point mutations

Given the surprising finding that exposure to well-known mutagens, despite the increase in damage, did not result in mtDNA mutations, we hypothesized that mitophagy may be working to remove the damaged mitochondria before having a chance to form mutations. To test this hypothesis, we conducted a similar MA exposure experiment in mitophagy deficient strains. We chose *dct-1* and *pink-1* deficient strains because *dct-1* and *pink-1* (BNIP-3 and PINK1 human homologs) are involved in two independent mitophagy pathways in in *C. elegans*. Mitophagy-deficient *dct-1* mutants did not accumulate significant mtDNA lesions compared to control after exposure to CdCl_2_ (0.21 lesions/10kb), but did after exposure to AfB_1_ (1.33 lesions/10kb, *P* < 0.01; two-way ANOVA, Tukey HSD) (Figure 4A). *pink-1* mutants accumulated significantly higher levels of mtDNA damage after exposure to CdCl_2_ and AfB_1_ respectively compared to control (1.03 and 0.72 lesions/10kb, *P* < 0.01; two-way ANOVA, Tukey HSD) (Figure 4A). Neither *dct-1* or *pink-1* mutants accumulated higher levels of mtDNA damage after exposure to CdCl_2_ compared to wild-type (two-way ANOVA). *dct-1* mutants did accumulate significantly higher levels of mtDNA damage compared to wild-type after exposure to AfB_1_ (*P* < 0.05; two-way ANOVA, Tukey HSD) (Figure 4A). *pink-1* mutants may accumulate higher levels of mtDNA damage compared to wild-type after AfB_1_ exposure; this trended towards, but did not reach, significance (*P =* 0.08; two-way ANOVA, Tukey HSD) (Figure 4A).

**Figure 4.**
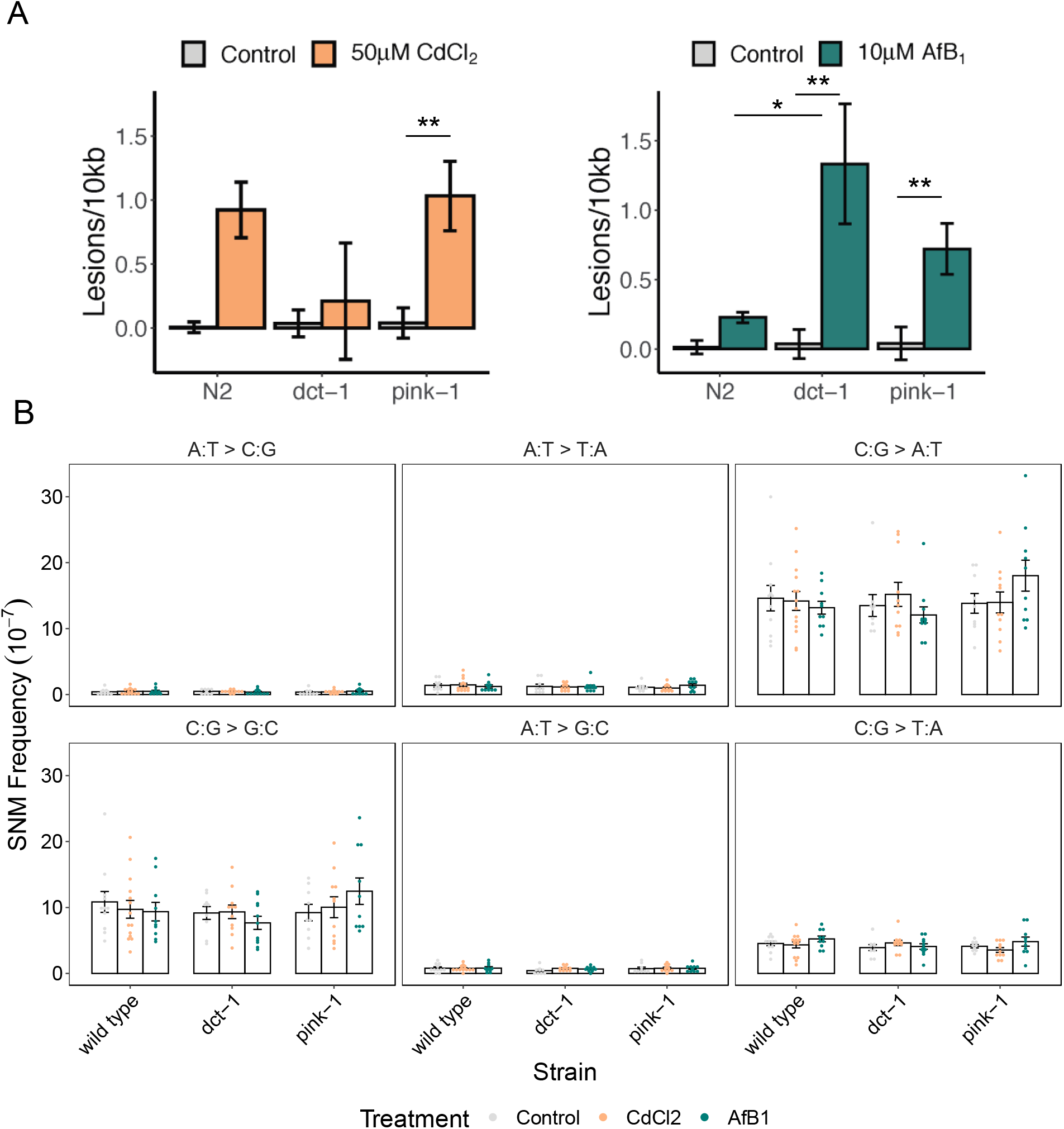
Mitophagy deficient mutants accumulate higher levels of mtDNA damage compared to wild-type, but exhibit no differences in mtDNA mutation frequencies after exposures. (A) mtDNA damage from pools of six individual age-synchronized L4 *C. elegans* after exposure to control (N = 8), 50µM CdCl_2_ (N = 8), or 10µM AfB_1_ (N = 8). *pink-1* mutants accumulated mtDNA damage after exposure to CdCl_2_, but not more than wild-type, while *dct-1* mutants did not accumulate mtDNA damage. Both mutants accumulated high levels of mtDNA damage after exposure to AfB_1,_ and *dct-1* accumulated significantly higher levels compared to wild-type exposed *C. elegans*. Error bars indicate standard error of the mean (* *P* < 0.05; ** *P* < 0.01; two-way ANOVA, Tukey HSD). (B) Mutation spectrum of control (gray), CdCl_2_ (gold), and AfB_1_ (green) treated MA lines in wild-type, *dct-1*, and *pink-1* strains. The mitophagy mutant *C. elegans* mtDNA mutational signatures were also dominated by C:G → A:T and C:G → G:C transversion mutations, and there was no effect of CdCl_2_ or AfB_1_ exposure on *dct-1* or *pink-1* mtDNA mutational signature (two-way ANOVA). Error bars indicate standard error of the mean.

We sequenced *dct-1* and *pink-1* MA lines after 50 generations of mutation accumulation in control, CdCl_2_, and AfB_1_-treated conditions. A total of 491, 727, and 574 total SNMs were detected after sequencing 11 control, 11 CdCl_2_, and 9 AfB_1_ *dct-1* MA lines, respectively. A total of 530, 696, and 752 total SNMs were detected after sequencing 9 control, 11 CdCl_2_, and 10 AfB_1_ *pink-1* MA lines, respectively. There was no effect of mitophagy deficiency or exposure on overall mtDNA mutation frequency (Supplementary Table S2, Supplementary Figure S8; two-way ANOVA, *P =* 0.3). The mutational spectrum of *dct-1* and *pink-1* was also dominated by C:G → A:T and C:G → G:C transitions, and C:G → T:A transversions (Figure 4B). There was no effect of CdCl_2_ or AfB_1_ on the mutational spectrum in either *dct-1* or *pink-1* mutants (two-way ANOVAs).

### Trinucleotide context mutagenesis

Mammalian mtDNA has been reported to have a distinctive mutational signature (60). Therefore, we analyzed the identity of the 5’ and 3’ neighboring nucleotides to investigate possible enrichment of SNMs in specific sequence contexts. Overall, the *C. elegans* mitochondrial genome does have a distinct trinucleotide mutational signature (Supplementary Figure S9). Specifically, C:G → T:A mutations occur in a G[C → T]C context, while C:G → A:T mutations occur mainly at A[C/G]A, X[C/G]T, T[C/G]A, and T[C/G]T sites. In order to determine the effect of chemical-induced mtDNA damage on trinucleotide context mutagenesis, we normalized the relative contribution of each mutation to the mean wild-type control contribution (Figure 5). In wild-type exposed to CdCl_2_, there were more C → G mutations in a C[C/G]A context compared to control (*P* < 0.05; Welch Two Sample T-test). There was also a trend of more T → C mutations in a C[T/A]T context compared to control (*P* = 0.06; one-way ANOVA, Welch Two Sample T-test). In AfB_1_ MA lines, we observed fewer C → G and C → T mutations in a C[C/G]C context, and more T → C mutations in a C[T/A]A context compared to control (*P* < 0.05; one-way ANOVA, Welch Two Sample T-test). In *dct-1* mutants, we observed more T → C mutations in CdCl_2_ MA lines in a C[T/A]A context compared to control (p < 0.05; one-way ANOVA, Welch Two Sample T-test), and more C → T in a A[C/G]T context and T → C in A[T/A]T and G[T/A]A contexts after exposure to AfB_1_ compared to control (*P* < 0.05; one-way ANOVA, Welch Two Sample T-test). In the *pink-1* mutants, we observed lower relative contributions of mutations at specific trinucleotide contexts compared to wild-type control, with the exception of more C → A mutations in a A[C/G]C context in AfB_1_ compared to *pink-1* control (*P* < 0.05; one-way ANOVA, Welch Two Sample T-test).

**Figure 5.**
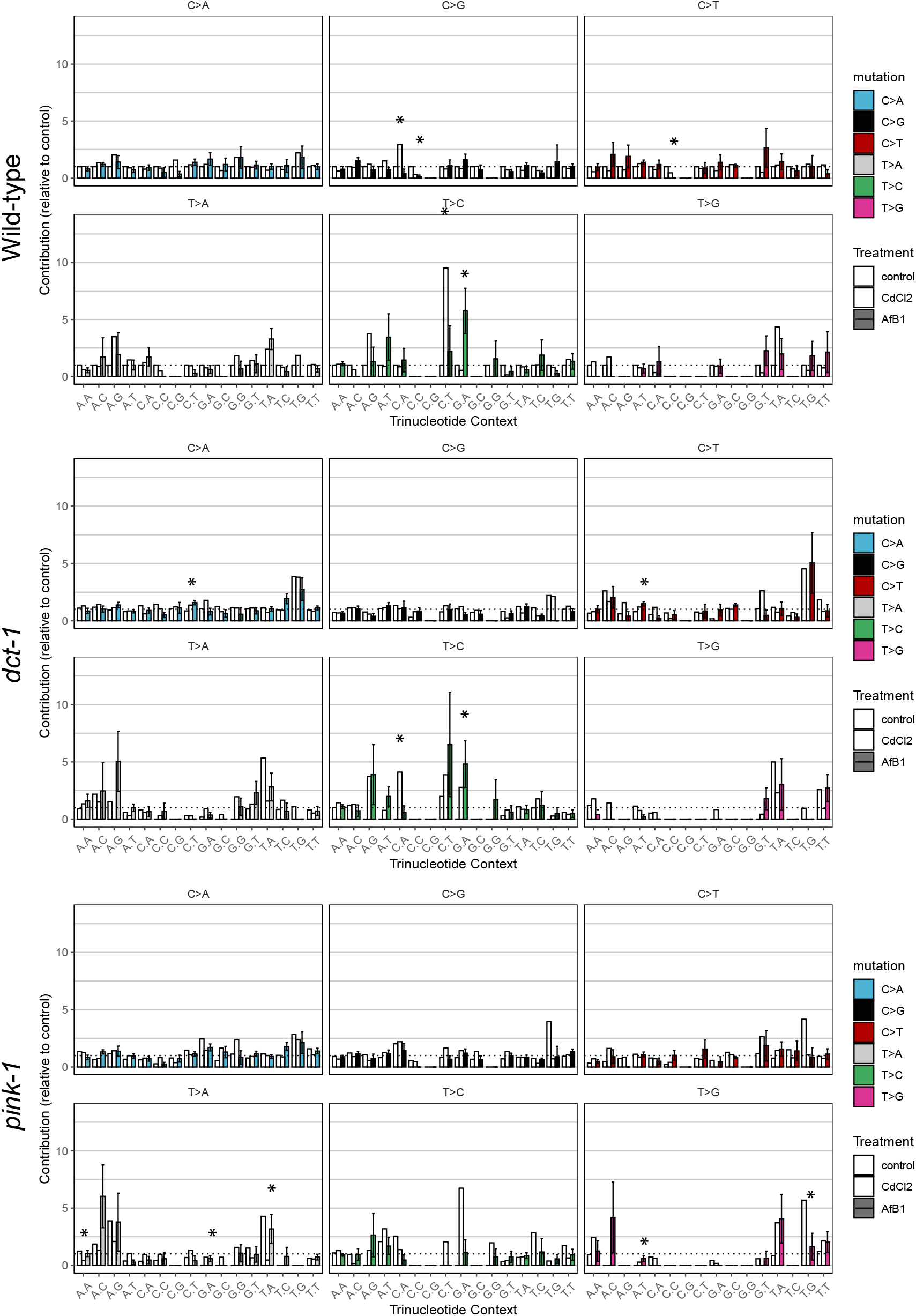
Trinucleotide context mutational signature of wild-type, *dct-1*, and *pink-1 C. elegans* after exposure to CdCl_2_ or AfB_1_. Contributions of each of the 96 possible trinucleotide mutations were determined for each MA line, and then normalized to the mean wild-type control contribution (show as a dashed line) to determine effects relative to wild-type control. Bar graphs show the relative mean and standard error of each trinucleotide context mutation, with each mutation represented by a different color and control, CdCl_2_, and AfB_1_ treatment as increasing color hues. Potential effects of each exposure compared to control were determined within each mutation type (Welch Two Sample T-test) only after a significant ANOVA was determined. * *P* < 0.05.

We used the software *MutationalPatterns* to determine the cosine similarities of trinucleotide context mutational signatures between strain and treatment. As there were very few effects in the enrichment of trinucleotide mutations due to treatment as described above, it was unsurprising that the trinucleotide context mutational signatures were highly similar between strains and treatment (Supplementary Figure S9). Indeed, the cosine similarity values across the three strains and three conditions were all close to 1 (Supplementary Table S3). We next determined the similarity of these mutational signatures to known mutational processes as described in the COSMIC database (Supplementary Table S4). It is critical to note that the COSMIC signatures are derived from somatic nuclear mutagenesis from human cancer genomes, and *C. elegans* germline mitochondrial mutational processes are likely very different. Taking this into consideration, it was striking that across all of our samples we detected high cosine similarity to the single base substitution (SBS) mutational signature that is associated with nucleotide excision repair deficiency (SBS24), which is absent in mitochondria. There was also high similarity with signatures associated with tobacco exposures (SBS4, SBS29). Tobacco products contains high levels of cadmium and smoke contains high levels of benzo[a]pyrene, which results in the formation of DNA adducts similar to those caused by AfB_1_. There also exists high similarity between signatures in which high levels of ROS is the proposed etiology (SBS18), in addition to defective base excision repair due to ROS-induced DNA damage (SBS36) and replication error across abasic sites (SBS13a). The similarities of *C. elegans* mtDNA signatures with certain COSMIC signatures are likely due to the high C:G → A:T mutational bias in the *C. elegans* mitochondrial genome. Cosine similarity values are located in Supplementary Table S4 and S5.

## DISCUSSION

Mutation accumulation experiments in *C. elegans* and other organisms have determined that chemical exposures contribute to mutagenesis in the nuclear genome (61, 62). However, the role of chemicals in mtDNA mutagenesis is not well studied (63). There is some evidence that mtDNA is resistant to single nucleotide mutations in somatic tissues after exposure to the polycyclic aromatic hydrocarbon benzo[a]pyrene and hydrogen peroxide (8, 14), and in germ cells after exposure to hydrogen peroxide (64). As mitophagy is responsible for degrading organelles with damaged mtDNA (22) and purifying selection against highly deleterious mutations in the germline (30), a widely accepted theory in the field is that mitophagy regulates mtDNA mutational processes. Indeed, in various models including a study in *C. elegans*, placing a Pol γ mutator strain in a mitophagy-deficient background resulted in increased mtDNA mutation frequency (65, 66). However, to our knowledge, empirical evidence investigating the role of mitophagy in ameliorating chemical-induced mtDNA damage, or mutations caused by damage, had not yet been investigated. Therefore, we conducted a mutation accumulation line experiment to determine the role of mitophagy in chemical-induced germline mtDNA mutagenesis (Figure 2). We found that *C. elegans* mtDNA is resistant to germline mutagenesis from exposure to known nuclear mutagens Cd and AfB_1_, despite accumulating high levels of mtDNA damage, extending the results observed previously with somatic mtDNA mutagenesis. Unexpectedly, we found that this resistance appears to be independent of mitophagy.

*C. elegans* continues to advance fundamental understanding of mitochondrial structure and function pertaining to human health and disease due to the conservation of mtDNA genes and various pathways (67). *C. elegans* is also a key research organism in the fields of evolutionary biology (32) and toxicology (68, 69), and has provided many insights into our knowledge of mtDNA mutagenesis and mitochondrial toxicity. For example, previous mutation accumulation line experiments with *C. elegans* have been instrumental in our understanding of mtDNA mutation rates (7, 33). However, various intricacies of mitochondrial genomics in *C. elegans* and other species render the study of mtDNA mutagenesis highly complex (70). The mitochondrial genome exists in multiple copies per organelle and cell, meaning that multiple genomes can pass through the germline bottleneck (∼60 copies in *C. elegans*) (33), potentially allowing selection to still act on the organelle and cellular level even in a neutral mutation accumulation design (71). Because mtDNA is polyploid, the frequency of each *de novo* single nucleotide mutation is often lower than the limit of detection of traditional next-generation sequencing (error rate of 1 in 10^3^ base pairs). Furthermore, mutations that result from DNA damage are likely even more rare than spontaneous mutagenesis (72). Therefore, we employed Duplex Sequencing in order to accurately detect *de novo* variants as low as 1 in 10^7^ bases, and hypothesized that inhibiting mitophagy would prevent purifying selection from acting within an individual during a mutation accumulation line experiment.

We sequenced 96 MA lines after about 50 generations of mutation accumulation each, resulting in a total of ∼4,800 generations of mutation accumulation. Overcoming the technical limitations of conventional next-generation sequencing by using Duplex Sequencing allowed us to detect a total of 6,040 SNMs, which is about 250-fold higher than any previous *C. elegans* mitochondrial mutation accumulation line experiment to date. The results of this Duplex Seq and MA approach were not biased by selection, as dN/dS rations were ∼1 across all MA lines (Supplementary Table S6), as has been a potential artifact and concern in previous studies (73, 74). Although proportionally similar across transition and transversion mutations, we observed roughly a 5-fold increase in mutation frequency compared to a recent study that conducted Duplex Sequencing to investigate mtDNA mutagenesis in wild-type *C. elegans*, which could possibly be attributed to the higher number of generation bottlenecks in our experiment (50 generations of one individual versus three generations of passaging 10 individuals) or other subtle differences in methodological and computational parameters (35). However, we did observe a mutation spectrum that is highly consistent with Waneka et al. (35). High frequencies of C:G → A:T and C:G → G:C transversion mutations support the conclusion that oxidative damage may drives mtDNA mutagenesis in *C. elegans*, though the mechanism underlying this spectrum remains an exciting area of future study. *C. elegans* appear to lack some homologs of *E. coli* 7,8-dihydro-8-oxoguanine (8-oxo-G) repair enzymes (MutY and MutM), as well as OGG1, which may explain the preponderance of transversion mutations (75–77). An earlier mutation accumulation study demonstrated that under extreme drift, the intracellular oxidizing environment increased in *C. elegans* and resulted in oxidative damage to the nuclear genome (78). This has not been measured in the mitochondrial genome, but could explain the high C:G → A:T mutation frequency in our MA experiment. We observed a dominant G[C → T]C trinucleotide mutation across all strains and treatments (Supplemental Figure S6). This signature has been detected in various studies including human mitochondrial studies in aged populations, and has also been observed in some cancers, suggesting that either deoxycytidine deamination readily occurs in the *C. elegans* mitochondrial genome, or oxidized cytosines contribute to this mutational pattern (79). Cytosine deamination due to oxidative stress results in an excess of C → T over G → A mutations on the mtDNA heavy strand in many different organisms including *Drosophila* (8), mouse (80), and human mtDNA (81), and is likely the primary source of Pol γ error-induced mtDNA mutagenesis (10). However, our results did not reveal any strand asymmetry in mutation frequencies in wild-type *C. elegans* as would be predicted, particularly since oxidative damage is the prevailing model of mtDNA mutagenesis in *C. elegans* (35). Lack of strand-asymmetric mutagenesis could be because the mechanism of mtDNA replication in *C. elegans* may deviate from the single-strand displacement theory of other organisms (82), perhaps restricting exposure of damage-prone single-stranded mtDNA to oxidative stress and thus leading to no observable strand bias. This remains an intriguing area of future study – especially the compensatory mechanisms that have evolved to evade high rates of oxidative damage-induced mutagenesis. An earlier *C. elegans* MA study determined that elevated ROS levels due to an electron transport chain complex I deficiency (*gas-1* mutant) resulted in a 3-fold increase in mtDNA copy number after almost 50 generations of mutation accumulation (34). Wernick *et al*. speculate that this is perhaps a compensatory mechanism to avoid the accumulation of potentially deleterious mtDNA variants when under extreme drift. It is important to note however that in the *gas-1* study, only five MA lines were sequenced after 50 generations and only five SNVs were detected among all five MA lines (three of which were already present in the *gas-1* progenitor). We did not investigate mtDNA copy number over generations, although it is possible that an increase in mtDNA replication under stress or other compensatory mechanisms contributed to our results (83, 84). However, the concentrations of CdCl_2_ and AfB_1_ at which the MA line experiments were conducted did not affect mtDNA copy number levels initially at G0 (Supplementary Figure S2). Alternatively, it is possible that high levels of endogenous oxidative damage and sequential spontaneous mutagenesis could overwhelm our ability to detect other forms of rare, damage-induced mutagenesis.

To our knowledge, no mutation accumulation experiment has investigated the effects of Cd exposure on mtDNA or in *C. elegans*. Cd is a ubiquitous heavy metal that is released from a range of sources, including, but not limited to, industrial processes such as mining and smelting. Combustion and wastewater contamination pollute terrestrial and aquatic environments and contaminate air, crops, and water. The most significant sources of human exposure to Cd are from contaminated food, smoking, e-waste sites, and even in products such as toys, jewelry, and plastics (85). Cd is a known carcinogen (86) and inhibits DNA repair mechanisms such as mismatch repair (37, 87–89), base excision repair (90), and many other zinc-dependent enzymes, and also induces oxidative stress, likely via antioxidant depletion mechanisms. Cd preferentially targets cellular components, including mitochondria (91). However, the genotoxic and mutagenic effects of Cd on mtDNA are unknown. Ultimately, this led us to investigate the effects of chronic, low-level exposure to CdCl_2_ on mtDNA mutagenesis in *C. elegans*.

To our surprise, we observed no effect of exposure to 50 µM CdCl_2_ on *C. elegans* mtDNA mutational spectrum across all genotypes, even though this exposure results in high levels of mtDNA damage (Figure 1A, Figure 4A). Overall, there was almost no effect on the trinucleotide context mutational signature after exposure to CdCl_2_, except for an enrichment of a C → G transversion mutation in a C[N]A context compared to control in wild-type *C. elegans*, which may be indicative of oxidative damage. 2,5-diamino-4H-imidazole-4-one (Iz) lesions contribute to C → G transversions in *E coli* (92). Intriguingly, a recent study determined that human Pol γ is more prone to preferentially misincorporate a G opposite 2,2,4-triamino-5(2 *H*)-oxazolone (Oz), the common hydrolyzed product of Iz (93, 94). Though we did not have the capability to identify the specific type of mtDNA lesions that are a result of Cd exposure (the long-amplification quantitative PCR assay used in this study cannot differentiate between different types of damage to mtDNA such as breaks, oxidized bases, or adducts), the preponderance of this C → G transversion mutation could suggest that Cd causes a variety of oxidation damage, possibly not just 8-oxo-7,8-dihydro-2′-deoxyguanosine (8-oxodG) lesions which are likely repaired in the mitochondria and contribute to C → A/G → T transversions. The mechanism is which oxidative damage and repair contributes to mtDNA mutational spectrum in *C. elegans* mtDNA mutagenesis is unclear, but there is evidence that *C. elegans* is able to repair at least some forms of oxidative mtDNA damage (77, 95–98), despite the aforementioned absence of clear homologues of some BER enzymes. Neither of the mitophagy mutant strains accumulated significantly higher levels of mtDNA damage after exposure to 50µM CdCl_2_ compared to exposed wild-type. This suggests that mitophagy is not the primary mechanism responsible for removal of damaged mtDNA or mtDNA mutations, at least at this level of mtDNA damage and frequency of mtDNA mutations. *dct-1* mutants do have an enrichment of one T → C transition mutation in a C[N]A context in Cd conditions compared to control, but the process underlying this mutation is unknown. It is possible that the quantity of lesions formed at this concentration of Cd was not enough to contribute to mtDNA mutagenesis. However, the internal body burden of Cd in *C. elegans* after exposure to 50µM CdCl_2_ (Supplementary Table S7) is equivalent to levels detected in human blood (86), suggesting that this concentration is relevant to human exposure.

This study contributes to evidence suggesting that *C. elegans* mtDNA mutagenesis may be driven by endogenous oxidative damage; however, as mentioned, *C. elegans* can repair at least some oxidative mtDNA damage. Therefore, to investigate the effect of irreparable mtDNA damage, we investigated mtDNA mutational spectrum and signatures after exposure to AfB_1_. AfB_1_ is a mycotoxin produced by the fungus *Aspergillus flavus*, and is one of the leading causes of hepatocellular carcinoma worldwide. *A. flavus* grows on staple grains and legumes, and when consumed, is metabolized to an epoxide that intercalates 5’ to guanine targets, resulting in stable AfB_1_-N^7^-G adducts that cause G → T mutations in the nuclear genome, primarily in CGC trinucleotide contexts (39). AfB_1_ exposure results in a similar mutational spectrum in the *C. elegans* nuclear genome that has been observed in human and mammalian studies (62, 99). However, to our knowledge, no study has investigated effects on mtDNA mutagenesis, even though it has been known for decades that AfB_1_ preferentially attacks mtDNA over nuclear DNA (42).

We measured very small effects of 10µM AfB_1_ on mtDNA mutational spectrum in wild-type or mitophagy-deficient *C. elegans* after 50 generations of exposure, even though AfB_1_ exposure resulted in significant levels of mtDNA damage which were exacerbated in the *dct-1* mitophagy mutant. We do observe a trend towards an increase in C:G → A:T transition mutations in AfB_1_-exposed *pink-1* mutants compared to wild-type (1.4-fold greater mutation frequency, *P =* 0.07, two-way ANOVA, Tukey HSD). This suggests that perhaps *pink-1* may play a role in ameliorating mtDNA damage or mediating purifying selection against deleterious mutations. In fact, we observe that over 50 generations of mutation accumulation, *pink-1* mutants have a greater fitness decline than wild-type and *dct-1* mutants, suggesting that they are accumulating more deleterious mutations (Supplementary Figure S10). Of course, we cannot rule out that this is due to mutations in the nuclear genome or nongenetic effects. However, it is not obvious why a mitophagy deficiency would elevate nuclear DNA mutagenesis, although it is conceivable that inhibiting mitophagy may attenuate mitochondria-nuclear DNA repair crosstalk mechanisms (100).

The trinucleotide context mutational signature in AfB_1_ MA lines does not reveal the clear signature that has been observed in nuclear genome studies. There is a significant but small enrichment of C → A mutations in a A[C/G]C context in AfB_1_ MA lines compared to control in *pink-1* mutants, which varies slightly from the known nuclear C[G]C context, suggesting a subtle but unique mechanism of aflatoxin damage-induced mutagenesis in the *C. elegans* mitochondrial genome. The unique AfB_1_ signature may result from the fact that damage removal processes are different for the two genomes; AfB_1_– induced damage in the nuclear genome is repaired by the enzyme NEIL1 as well as NER (101), but NER does not operate in the mitochondria and *C. elegans* does not appear to have a NEIL1 gene.

The lack of an effect of mitophagy deficiency on mutation accumulation in these studies appears initially to be at odds with reports of a role for mitophagy in regulating selection against inheritance of mtDNA mutations in germ cells. A few possibilities may explain this. First, mitophagy may work efficiently only in the case of high-frequency or highly-deleterious mutations. It is possible that *de novo* mutations do not have functional consequences on organelle function, and therefore evade targeted degradation via mitophagy. Second, it is possible that the levels of chemical-induced mtDNA damage may not be high enough to contribute significantly to mtDNA mutagenesis in this study. We consider this to be unlikely however, as we used an exposure concentration of AfB_1_ that is considerably higher than a previous *C. elegans* study that detected nuclear mutational signatures after exposure to AfB_1_ at concentrations 33 and three-fold lower than this study (0.3µM and 3µM AfB_1_). Volkova *et al*. detected a clear mutational signature that shared high similarity to the AfB_1_ mutational signature observed in humans (C → A/G → T mutations) after only one generation of exposure (62, 99). Because we conducted targeted capture sequencing of just mtDNA, we were unable to analyze effects of exposure on nuclear genome mutagenesis. However, given those results, it is likely that a significant amount of *de novo* variants arose in the nuclear genome in our MA study after 50 generations of exposure to a higher dose of AfB_1_. Third, with Duplex Sequencing, we were only able to measure single nucleotide point mutations, not large insertions or deletions. Recent work indicates that mtDNA in somatic cells accumulates a large number of individually uncommon deletions with age, likely as a result of polymerase error (102). It is possible that chemical-induced adducts (in the case of AfB_1_) or oxidative damage (CdCl_2_) may exacerbate polymerase error-mediated deletion accumulation.

Overall, our study suggests that in *C. elegans*, mitochondria harbor important quality control processes that are perhaps complementary or redundant to the two mitophagy pathways investigated in this study, resulting in a remarkable resistance to chemical-induced mutagenesis. A recent study in *Drosophila* found that unhealthy organelles failed to import nuclear-encoded factors essential for mtDNA replication and biogenesis. This resulted in the replication of wild-type mtDNA in healthy organelles, and not mutant mtDNA (103). Perhaps a “replication-competition” model remains to be discovered in *C. elegans*. Direct degradation of damaged or mutant mtDNA may also play a protective role (104), though to our knowledge, *C. elegans* lacks a homolog for mitochondrial genome maintenance exonuclease (MGME-1) that degrades linear and damaged mtDNA at the mtDNA replisome. Mechanisms that target paternal mitochondria and mtDNA for degradation in *C. elegans* are known, and an intriguing possibility is that similar pathways that target damaged or mutant mtDNA are present the germline, which could suppress accumulation of deleterious mtDNA variants (105, 106). On the organelle level, mitochondrial fusion and fission play a significant role in mediating mitochondrial quality control and are prerequisites for mitophagy (20, 23). At 50µM CdCl_2_ and 10µM AfB_1_ exposure, we did not observe fragmentation of the *C. elegans* body wall muscle mitochondrial network in G0 wild-type *C. elegans* compared to control. We observed increased mitochondrial fusion at lower levels of Cd and AfB_1_exposure, with recovery at higher doses, which suggests that mitochondrial dynamics are responding to Cd and AfB_1_-induced toxicity (Supplementary Figure S5). Additionally, we do not observe any effects on mitochondrial respiration in all three strains, so it is not likely that these exposures severely compromised mitochondrial function (Supplementary Figure S6). However, we propose that future studies investigate the role of mitochondrial dynamics in organelle and mtDNA turnover and biogenesis in the context of other chemical exposures. Lastly, another potential mechanism of ameliorating the removal of damaged mtDNA in the germline on the organismal level is germline apoptosis, which has previously been reported to increase after Cd and AfB_1_ exposure in *C. elegans*, though not at the concentrations that were used in our study (107, 108).

The use of chemicals in society has resulted in over 86,000 chemicals of unknown toxicity in production (109), in addition to the hundreds of thousands of tons of pharmaceutical and industrial waste, increases in air pollution, and other sources of pollution that impact the health of hundreds of millions of people daily around the globe (110). Many chemicals can cause genome instability, which drives cancer development and other diseases; therefore it is necessary to better understand how low-level exposures to environmental agents can promote mutagenesis (111, 112). mtDNA mutagenesis contributes to cancer and other diseases (60, 113–116), yet is far less understood than nuclear DNA mutagenesis. Mitochondria are vital organelles that are highly susceptible to toxicity and genomic damage (16, 117, 118). MtDNA mutations accumulate with age, yet there remains conflicting evidence in the literature for the role of genotoxicant exposure in mtDNA mutagenesis (63). Surprisingly, we found that in *C. elegans*, after thousands of generations of continuous chemical exposure, with thousands of SNMs detected, and in the context of loss of two key mitophagy genes, mtDNA is resistant to damage-induced single nucleotide mutations. Future studies to investigate the effect of more chemicals in various genetic backgrounds may elucidate mechanisms in which mitochondria resist chemical-induced point mutations.

## AVAILABILITY

The Duplex-Seq-Pipeline is written in Python and R, but has dependencies written in other languages. The Duplex-Seq-Pipeline software has been tested to run on Linux, Windows WSL1, Windows WSL2 and Apple OSX. The software can be obtained at https://github.com/Kennedy-Lab-UW/Duplex-Seq-Pipeline under the BSD license.

## ACCESSION NUMBERS

The data are available as raw reads under SRA Accession SRP350474 (BioProject PRJNA787252).

## SUPPLEMENTARY DATA

Supplementary Data are available at NAR online.

## ACKNOWLEDGEMENT

We would like to thank members of the Duke Center for Genomics and Computational Biology, particularly Dr. Nicolas Devos and Nicholas Hoang for their help with library preparation and sequencing. We thank Inigo Martincorena for his help with the *dNdScv* package. We would also like to thank members of the Meyer, Baugh, Wernegreen, and Jayasundara labs at Duke University for their thoughtful feedback throughout the design, implementation, and analysis stages of this project, and feedback on this manuscript.

## FUNDING

This work was supported by the National Institutes of Health [(F31 ES030588 to TCL, P42ES010356 to JNM, IRM supported by T32ES021432]; Triangle Center for Evolutionary Medicine (TriCEM) Graduate Student Award to TCL; Duke University School of Medicine Sequencing and Genomics Technologies Core Facility Voucher to TCL and JNM; Congressionally Directed Medical Research Programs [W81XWH-16-1-0579, W81XWH-18-1-0339]; National Human Genome Research Institute [R21 HG011229] to SRK. Funding for open access charge: National Institutes of Health.

## CONFLICT OF INTEREST

S.R.K. is an equity holder and paid consultant for Twinstrand Biosciences, a for-profit company commercializing Duplex Sequencing. No Twinstrand products were used in the generation of the data.

## REFERENCES

1. Bannwarth, S., Procaccio, V., Lebre, A.S., Jardel, C., Chaussenot, A., Hoarau, C., Maoulida, H., Charrier, N., Gai, X., Xie, H.M., et al. (2013) Prevalence of rare mitochondrial DNA mutations in mitochondrial disorders. J. Med. Genet., 50, 704–14.

2. Hertweck, K.L. and Dasgupta, S. (2017) The Landscape of mtDNA Modifications in Cancer: A Tale of Two Cities. Front. Oncol., 7, 262.

3. Ju, Y.S. eo., Alexandrov, L.B., Gerstung, M., Martincorena, I., Nik-Zainal, S., Ramakrishna, M., Davies, H.R., Papaemmanuil, E., Gundem, G., Shlien, A., et al. (2014) Origins and functional consequences of somatic mitochondrial DNA mutations in human cancer. Elife, 3, 1–28.

4. Tuppen, H.A.L., Blakely, E.L., Turnbull, D.M. and Taylor, R.W. (2009) Mitochondrial DNA mutations and human disease. BBA - Bioenerg., 1797, 113–128.

5. Brown, W.M., George, M. and Wilson, A.C. (1979) Rapid evolution of animal mitochondrial DNA. Proc. Natl. Acad. Sci. U. S. A., 76, 1967–71.

6. Brown, W.M., Prager, E.M., Wang, A. and Wilson, A.C. (1982) Mitochondrial DNA sequences of primates: Tempo and mode of evolution. J. Mol. Evol., 18, 225–239s.

7. Denver, D.R., Morris, K., Lynch, M., Vassilieva, L.L. and Thomas, W.K. (2000) High Direct Estimate of the Mutation Rate in the Mitochondrial Genome of Caenorhabditis elegans. Science (80-.)., 289, 2342–2345.

8. Itsara, L.S., Kennedy, S.R., Fox, E.J., Yu, S., Hewitt, J.J., Sanchez-Contreras, M., Cardozo-Pelaez, F. and Pallanck, L.J. (2014) Oxidative stress is not a major contributor to somatic mitochondrial DNA mutations. PLoS Genet., 10, e1003974.

9. Kennedy, S.R., Salk, J.J., Schmitt, M.W. and Loeb, L.A. (2013) Ultra-sensitive sequencing reveals an age-related increase in somatic mitochondrial mutations that are inconsistent with oxidative damage. PLoS Genet., 9, e1003794.

10. Sanchez-Contreras, M., Sweetwyne, M.T., Kohrn, B.F., Tsantilas, K.A., Fredrickson, J., Whitson, J.A., Campbell, M.D., Rabinovitch, P.S., Marcinek, D.J. and Kennedy, S.R. (2021) A mutational gradient drives somatic mutation accumulation in mitochondrial DNA and influences germline polymorphisms and genome composition. Nucleic Acids Res.

11. Krasich, R. and Copeland, W.C. (2017) DNA polymerases in the mitochondria: A critical review of the evidence. Front. Biosci. (Landmark Ed., 22, 692–709.

12. Cline, S.D. (2012) Mitochondrial DNA damage and its consequences for mitochondrial gene expression. Biochim. Biophys. Acta - Gene Regul. Mech., 1819, 979–991.

13. Gustafson, M.A., Sullivan, E.D. and Copeland, W.C. (2020) Consequences of compromised mitochondrial genome integrity. DNA Repair (Amst)., 93, 102916.

14. Valente, W.J., Ericson, N.G., Long, A.S., White, P.A., Marchetti, F. and Bielas, J.H. (2016) Mitochondrial DNA exhibits resistance to induced point and deletion mutations. Nucleic Acids Res., 44, 8513–8524.

15. Meyer, J.N., Leung, M.C.K., Rooney, J.P., Sendoel, A., Hengartner, M.O., Kisby, G.E. and Bess, A.S. (2013) Mitochondria as a target of environmental toxicants. Toxicol. Sci., 134, 1–17.

16. Meyer, J.N., Hartman, J.H. and Mello, D.F. (2018) Mitochondrial Toxicity. Toxicol. Sci., 162, 15–23.

17. Roubicek, D.A. and de Souza-Pinto, N.C. (2017) Mitochondria and mitochondrial DNA as relevant targets for environmental contaminants. Toxicology, 391, 100–108.

18. Nunnari, J. and Suomalainen, A. (2012) Mitochondria: In Sickness and in Health. Cell, 148, 1145–1159.

19. Meyer, J.N., Leuthner, T.C. and Luz, A.L. (2017) Mitochondrial fusion, fission, and mitochondrial toxicity. Toxicology, 391, 42–53.

20. Luz, A.L., Godebo, T.R., Smith, L.L., Leuthner, T.C., Maurer, L.L. and Meyer, J.N. (2017) Deficiencies in mitochondrial dynamics sensitize Caenorhabditis elegans to arsenite and other mitochondrial toxicants by reducing mitochondrial adaptability. Toxicology, 387, 81–94.

21. Luz, A.L., Rooney, J.P., Kubik, L.L., Gonzalez, C.P., Song, D.H. and Meyer, J.N. (2015) Mitochondrial Morphology and Fundamental Parameters of the Mitochondrial Respiratory Chain Are Altered in Caenorhabditis elegans Strains Deficient in Mitochondrial Dynamics and Homeostasis Processes. PLoS One, 10, e0130940.

22. Bess, A.S., Crocker, T.L., Ryde, I.T. and Meyer, J.N. (2012) Mitochondrial dynamics and autophagy aid in removal of persistent mitochondrial DNA damage in Caenorhabditis elegans. Nucleic Acids Res., 40, 7916–7931.

23. Bess, A.S., Leung, M.C.K., Ryde, I.T., Rooney, J.P., Hinton, D.E. and Meyer, J.N. (2013) Effects of mutations in mitochondrial dynamics-related genes on the mitochondrial response to ultraviolet C radiation in developing Caenorhabditis elegans. Worm, 2, e23763.

24. Jansen, R.P.S. and De Boer, K. (1998) The bottleneck: Mitochondrial imperatives in oogenesis and ovarian follicular fate. Mol. Cell. Endocrinol., 145, 81–88.

25. Bergstrom, C.T. and Pritchard, J. (1998) Germline bottlenecks and the evolutionary maintenance of mitochondrial genomes. Genetics, 149, 2135–2146.

26. Cree, L.M., Samuels, D.C., de Sousa Lopes, S.C., Rajasimha, H.K., Wonnapinij, P., Mann, J.R., Dahl, H.-H.M. and Chinnery, P.F. (2008) A reduction of mitochondrial DNA molecules during embryogenesis explains the rapid segregation of genotypes. Nat. Genet., 40, 249–254.

27. Cao, L., Shitara, H., Sugimoto, M., Hayashi, J.-I., Abe, K. and Yonekawa, H. (2009) New evidence confirms that the mitochondrial bottleneck is generated without reduction of mitochondrial DNA content in early primordial germ cells of mice. PLoS Genet., 5, e1000756.

28. Wai, T., Teoli, D. and Shoubridge, E.A. (2008) The mitochondrial DNA genetic bottleneck results from replication of a subpopulation of genomes. Nat. Genet., 40, 1484–1488.

29. Mishra, P. and Chan, D.C. (2014) Mitochondrial dynamics and inheritance during cell division, development and disease. Nat Rev Mol Cell Biol, 15, 634–646.

30. Valenci, I., Yonai, L., Bar-Yaacov, D., Mishmar, D. and Ben-Zvi, A. (2015) Parkin modulates heteroplasmy of truncated mtDNA in Caenorhabditis elegans. Mitochondrion, 20, 64–70.

31. Haroon, S., Li, A., Weinert, J.L., Fritsch, C., Ericson, N.G., Alexander-Floyd, J., Braeckman, B.P., Haynes, C.M., Bielas, J.H., Gidalevitz, T., et al. (2018) Multiple Molecular Mechanisms Rescue mtDNA Disease in C. elegans. Cell Rep., 22, 3115–3125.

32. Katju, V. and Bergthorsson, U. (2019) Old trade, new tricks: Insights into the spontaneous mutation process from the partnering of classical mutation accumulation experiments with high-throughput genomic approaches. Genome Biol. Evol., 11, 136–165.

33. Konrad, A., Thompson, O., Waterston, R.H., Moerman, D.G., Keightley, P.D., Bergthorsson, U. and Katju, V. (2017) Mitochondrial mutation rate, spectrum and heteroplasmy in Caenorhabditis elegans spontaneous mutation accumulation lines of differing population size. Mol. Biol. Evol., 34, 1319–1334.

34. Wernick, R.I., Estes, S., Howe, D.K. and Denver, D.R. (2016) Paths of Heritable Mitochondrial DNA Mutation and Heteroplasmy in Reference and gas-1 Strains of Caenorhabditis elegans. Front. Genet., 7.

35. Waneka, G., Svendsen, J.M., Havird, J.C. and Sloan, D.B. (2021) Mitochondrial mutations in Caenorhabditis elegans show signatures of oxidative damage and an AT-bias. Genetics, 219.

36. Schmitt, M.W., Kennedy, S.R., Salk, J.J., Fox, E.J., Hiatt, J.B. and Loeb, L.A. (2012) Detection of ultra-rare mutations by next-generation sequencing. Proc. Natl. Acad. Sci. U. S. A., 109, 14508–13.

37. Jin, Y.H., Clark, A.B., Slebos, R.J.C., Al-Refai, H., Taylor, J.A., Kunkel, T.A., Resnick, M.A. and Gordenin, D.A. (2003) Cadmium is a mutagen that acts by inhibiting mismatch repair. Nat. Genet., 34, 326–9.

38. Liu, J., Qu, W. and Kadiiska, M.B. (2009) Role of oxidative stress in cadmium toxicity and carcinogenesis. Toxicol. Appl. Pharmacol., 238, 209–14.

39. Chawanthayatham, S., Valentine, C.C., Fedeles, B.I., Fox, E.J., Loeb, L.A., Levine, S.S., Slocum, S.L., Wogan, G.N., Croy, R.G. and Essigmann, J.M. (2017) Mutational spectra of aflatoxin B1 in vivo establish biomarkers of exposure for human hepatocellular carcinoma. Proc. Natl. Acad. Sci. U. S. A., 114, E3101–E3109.

40. González-Hunt, C.P., Leung, M.C.K., Bodhicharla, R.K., McKeever, M.G., Arrant, A.E., Margillo, K.M., Ryde, I.T., Cyr, D.D., Kosmaczewski, S.G., Hammarlund, M., et al. (2014) Exposure to Mitochondrial Genotoxins and Dopaminergic Neurodegeneration in Caenorhabditis elegans. PLoS One, 9, e114459.

41. Leung, M.C.K., Goldstone, J. V, Boyd, W.A., Freedman, J.H. and Meyer, J.N. (2010) Caenorhabditis elegans generates biologically relevant levels of genotoxic metabolites from aflatoxin B1 but not benzo[a]pyrene in vivo. Toxicol. Sci., 118, 444–53.

42. Niranjan, A.B.G., Bhat, N.K. and Avadhani, N.G. (1982) Preferential Attack of Mitochondrial DNA by Aflatoxin B1 during Hepatocarcinogenesis. 215, 73–75.

43. Palikaras, K., Lionaki, E. and Tavernarakis, N. (2015) Coordination of mitophagy and mitochondrial biogenesis during ageing in C. elegans. Nature, 521, 525–528.

44. Brenner, S. (1974) The Genetics of Caenorhabditis elegans. Genetics, 77, 71–94.

45. Williams, P.L. and Dusenbery, D.B. (1988) Using the nematode caenorhabditis elegans to predict mammalian acute lethality to metallic salts. Toxicol. Ind. Health, 4, 469–478.

46. Boyd, W.A., McBride, S.J., Rice, J.R., Snyder, D.W. and Freedman, J.H. (2010) A high-throughput method for assessing chemical toxicity using a Caenorhabditis elegans reproduction assay. Toxicol. Appl. Pharmacol., 245, 153–9.

47. Cothren, S., Meyer, J. and Hartman, J. (2018) Blinded Visual Scoring of Images Using the Freely-available Software Blinder. Bio-Protocol, 8, 1–10.

48. Hartman, J.H., Smith, L.L., Gordon, K.L., Laranjeiro, R., Driscoll, M., Sherwood, D.R. and Meyer, J.N. (2018) Swimming Exercise and Transient Food Deprivation in Caenorhabditis elegans Promote Mitochondrial Maintenance and Protect Against Chemical-Induced Mitotoxicity. Sci. Rep., 8, 1–16.

49. Luz, A.L., Smith, L.L., Rooney, J.P. and Meyer, J.N. (2015) Seahorse Xfe 24 Extracellular Flux Analyzer-Based Analysis of Cellular Respiration in Caenorhabditis elegans. In Current protocols in toxicology. John Wiley & Sons, Inc., Hoboken, NJ, USA, Vol. 66, pp. 25.7.1–25.7.15.

50. Lant, B. and Derry, W.B. (2014) Fluorescent visualization of germline apoptosis in living Caenorhabditis elegans. Cold Spring Harb. Protoc., 2014, 420–427.

51. Moore, B.T., Jordan, J.M. and Baugh, L.R. (2013) WormSizer: High-throughput Analysis of Nematode Size and Shape. PLoS One, 8, 1–13.

52. Hodgkin, J. and Barnes, T.A. (1991) More is not better: brood size and population growth in a self-fertilizing nematode. Proc. R. Soc. London. Ser. B Biol. Sci., 246, 19–24.

53. Gonzalez-Hunt, C.P., Rooney, J.P., Ryde, I.T., Anbalagan, C., Joglekar, R. and Meyer, J.N. (2016) PCR-Based Analysis of Mitochondrial DNA Copy Number, Mitochondrial DNA Damage, and Nuclear DNA Damage. Curr. Protoc. Toxicol., 67, 20.11.1–20.11.25.

54. Leuthner, T.C., Hartman, J.H., Ryde, I.T. and Meyer, J.N. (2021) PCR-Based Determination of Mitochondrial DNA Copy Number in Multiple Species. Methods Mol. Biol., 2310, 91–111.

55. Denver, D.R., Feinberg, S., Estes, S., Thomas, W.K. and Lynch, M. (2005) Mutation rates, spectra and hotspots in mismatch repair-deficient Caenorhabditis elegans. Genetics, 170, 107–113.

56. Kennedy, S.R., Schmitt, M.W., Fox, E.J., Kohrn, B.F., Salk, J.J., Ahn, E.H., Prindle, M.J., Kuong, K.J., Shen, J.C., Risques, R.A., et al. (2014) Detecting ultralow-frequency mutations by Duplex Sequencing. Nat. Protoc., 9, 2586–2606.

57. Hoekstra, J.G., Hipp, M.J., Montine, T.J. and Kennedy, S.R. (2016) Mitochondrial DNA mutations increase in early stage Alzheimer disease and are inconsistent with oxidative damage. Ann. Neurol., 80, 301–306.

58. Blokzijl, F., Janssen, R., van Boxtel, R. and Cuppen, E. (2018) MutationalPatterns: Comprehensive genome-wide analysis of mutational processes. Genome Med., 10, 1–11.

59. Martincorena, I., Raine, K.M., Gerstung, M., Dawson, K.J., Haase, K., Van Loo, P., Davies, H., Stratton, M.R. and Campbell, P.J. (2017) Universal Patterns of Selection in Cancer and Somatic Tissues. Cell, 171, 1029–1041.e21.

60. Ju, Y.S. eo., Alexandrov, L.B., Gerstung, M., Martincorena, I., Nik-Zainal, S., Ramakrishna, M., Davies, H.R., Papaemmanuil, E., Gundem, G., Shlien, A., et al. (2014) Origins and functional consequences of somatic mitochondrial DNA mutations in human cancer. Elife, 3, 1–29.

61. Keith, N., Jackson, C.E., Glaholt, S.P., Young, K., Lynch, M. and Shaw, J.R. (2021) Genome-Wide Analysis of Cadmium-Induced, Germline Mutations in a Long-Term Daphnia pulex Mutation-Accumulation Experiment. Environ. Health Perspect., 129, 1–10.

62. Volkova, N. V., Meier, B., González-Huici, V., Bertolini, S., Gonzalez, S., Vöhringer, H., Abascal, F., Martincorena, I., Campbell, P.J., Gartner, A., et al. (2020) Mutational signatures are jointly shaped by DNA damage and repair. Nat. Commun., 11, 1–15.

63. Leuthner, T.C. and Meyer, J.N. (2021) Mitochondrial DNA Mutagenesis: Feature of and Biomarker for Environmental Exposures and Aging. Curr. Environ. Heal. Reports, 10.1007/s40572-021-00329-1.

64. Kauppila, J.H.K., Bonekamp, N.A., Mourier, A., Isokallio, M.A., Just, A., Kauppila, T.E.S., Stewart, J.B. and Larsson, N.G. (2018) Base-excision repair deficiency alone or combined with increased oxidative stress does not increase mtDNA point mutations in mice. Nucleic Acids Res., 46, 6642–6649.

65. Pickrell, A.M., Huang, C.H., Kennedy, S.R., Ordureau, A., Sideris, D.P., Hoekstra, J.G., Harper, J.W. and Youle, R.J. (2015) Endogenous Parkin Preserves Dopaminergic Substantia Nigral Neurons following Mitochondrial DNA Mutagenic Stress. Neuron, 87, 371–381.

66. Haroon, S., Li, A., Weinert, J.L., Fritsch, C., Ericson, N.G., Alexander-Floyd, J., Braeckman, B.P., Haynes, C.M., Bielas, J.H., Gidalevitz, T., et al. (2018) Multiple Molecular Mechanisms Rescue mtDNA Disease in C. elegans. Cell Rep., 22, 3115–3125.

67. Van Der Bliek, A.M., Sedensky, M.M. and Morgan, P.G. (2017) Cell biology of the mitochondrion. Genetics, 207, 843–871.

68. Leung, M.C.K., Williams, P.L., Benedetto, A., Au, C., Helmcke, K.J., Aschner, M. and Meyer, J.N. (2008) Caenorhabditis elegans: An Emerging Model in Biomedical and Environmental Toxicology. Toxicol. Sci., 106, 5–28.

69. Hunt, P.R. (2016) The C. elegans model in toxicity testing. J. Appl. Toxicol., 10.1002/jat.3357.

70. Schaack, S., Ho, E.K.H. and Macrae, F. (2020) Disentangling the intertwined roles of mutation, selection and drift in the mitochondrial genome. Philos. Trans. R. Soc. B Biol. Sci., 375, 20190173.

71. Stewart, J.B., Freyer, C., Elson, J.L. and Larsson, N.-G. (2008) Purifying selection of mtDNA and its implications for understanding evolution and mitochondrial disease. Nat. Rev. Genet., 9, 657–662.

72. Salk, J.J. and Kennedy, S.R. (2020) Next-Generation Genotoxicology: Using Modern Sequencing Technologies to Assess Somatic Mutagenesis and Cancer Risk. Environ. Mol. Mutagen., 61, 135–151.

73. Schaack, S., Ho, E.K.H. and Macrae, F. (2020) Disentangling the intertwined roles of mutation, selection and drift in the mitochondrial genome. Philos. Trans. R. Soc. B Biol. Sci., 375, 20190173.

74. Samstag, C.L., Hoekstra, J.G., Huang, C.H., Chaisson, M.J., Youle, R.J., Kennedy, S.R. and Pallanck, L.J. (2018) Deleterious mitochondrial DNA point mutations are overrepresented in Drosophila expressing a proofreading-defective DNA polymerase γ. PLoS Genet., 14, 1–27.

75. Jansson, K., Blomberg, A., Sunnerhagen, P. and Alm Rosenblad, M. (2010) Evolutionary loss of 8-oxo-G repair components among eukaryotes. Genome Integr., 1, 1–10.

76. Arczewska, K.D., Baumeier, C., Kassahun, H., SenGupta, T., Bjørås, M., Kuśmierek, J.T. and Nilsen, H. (2011) Caenorhabditis elegans NDX-4 is a MutT-type enzyme that contributes to genomic stability. DNA Repair (Amst)., 10, 176–187.

77. Sanada, U., Yonekura, S.I., Kikuchi, M., Hashiguchi, K., Nakamura, N., Yonei, S. and Zhang-Akiyama, Q.M. (2011) NDX-1 protein hydrolyzes 8-oxo-7, 8-dihydrodeoxyguanosine-5′-diphosphate to sanitize oxidized nucleotides and prevent oxidative stress in Caenorhabditis elegans. J. Biochem., 150, 649–657.

78. Christy, S.F., Wernick, R.I., Lue, M.J., Velasco, G., Howe, D.K., Denver, D.R. and Estes, S. (2017) Adaptive Evolution under Extreme Genetic Drift in Oxidatively Stressed Caenorhabditis elegans. Genome Biol. Evol., 9, 3008–3022.

79. Degtyareva, N.P., Saini, N., Sterling, J.F., Placentra, V.C., Klimczak, L.J., Gordenin, D.A. and Doetsch, P.W. (2019) Mutational signatures of redox stress in yeast single-strand DNA and of aging in human mitochondrial DNA share a common feature. PLoS Biol., 17, 1–28.

80. Arbeithuber, B., Hester, J., Cremona, M.A., Stoler, N., Zaidi, A., Higgins, B., Anthony, K., Chiaromonte, F., Diaz, J. and Id, K.D.M. (2020) Age-related accumulation of de novo mitochondrial mutations in mammalian oocytes and somatic tissues. PLoS Biol., 18, e3000745.

81. Kennedy, S.R., Salk, J.J., Schmitt, M.W. and Loeb, L. a. (2013) Ultra-Sensitive Sequencing Reveals an Age-Related Increase in Somatic Mitochondrial Mutations That Are Inconsistent with Oxidative Damage. PLoS Genet., 9, e1003794.

82. Lewis, S.C., Joers, P., Willcox, S., Griffith, J.D., Jacobs, H.T. and Hyman, B.C. (2015) A rolling circle replication mechanism produces multimeric lariats of mitochondrial DNA in Caenorhabditis elegans. PLoS Genet., 11, e1004985.

83. Gitschlag, B.L., Kirby, C.S., Samuels, D.C., Gangula, R.D., Mallal, S.A. and Patel, M.R. (2016) Homeostatic Responses Regulate Selfish Mitochondrial Genome Dynamics in C.??elegans. Cell Metab., 24, 91–103.

84. Lin, Y.-F., Schulz, A.M., Pellegrino, M.W., Lu, Y., Shaham, S. and Haynes, C.M. (2016) Maintenance and propagation of a deleterious mitochondrial genome by the mitochondrial unfolded protein response. Nature, 533, 416–9.

85. World Health Organization (2010) Preventing disease through healthy environments. Exposure to cadmium: a major public health concern. World Heal. Organ.

86. Agency for Toxic Substances and Disease Registry (2019) Toxicological Profile for Cadmium Atlanta, GA.

87. Hsu, T., Tsai, H.-T., Huang, K.-M., Luan, M.-C. and Hsieh, C.-R. (2010) Sublethal levels of cadmium down-regulate the gene expression of DNA mismatch recognition protein MutS homolog 6 (MSH6) in zebrafish (Danio rerio) embryos. Chemosphere, 81, 748–754.

88. Abbà, S., Vallino, M., Daghino, S., Di Vietro, L., Borriello, R. and Perotto, S. (2011) A PLAC8-containing protein from an endomycorrhizal fungus confers cadmium resistance to yeast cells by interacting with Mlh3p. Nucleic Acids Res., 39, 7548–7563.

89. Sherrer, S.M., Penland, E. and Modrich, P. (2018) The mutagen and carcinogen cadmium is a highaffinity inhibitor of the zinc-dependent MutLα endonuclease. Proc. Natl. Acad. Sci. U. S. A., 115, 7314–7319.

90. Antoniali, G., Marcuzzi, F., Casarano, E. and Tell, G. (2015) Cadmium treatment suppresses DNA polymerase δ catalytic subunit gene expression by acting on the p53 and Sp1 regulatory axis. DNA Repair (Amst)., 35, 90–105.

91. Lee, W.K. and Thévenod, F. (2020) Cell organelles as targets of mammalian cadmium toxicity Springer Berlin Heidelberg.

92. Neeley, W.L., Delaney, J.C., Henderson, P.T. and Essigmann, J.M. (2004) In vivo bypass efficiencies and mutational signatures of the guanine oxidation products 2-aminoimidazolone and 5-guanidino-4-nitroimidazole. J. Biol. Chem., 279, 43568–43573.

93. Kino, K., Kawada, T., Hirao-Suzuki, M., Morikawa, M. and Miyazawa, H. (2020) Products of oxidative guanine damage form base pairs with guanine. Int. J. Mol. Sci., 21, 1–21.

94. Kino, K., Sugasawa, K., Mizuno, T., Bando, T., Sugiyama, H., Akita, M., Miyazawa, H. and Hanaoka, F. (2009) Eukaryotic DNA polymerases α, β and ε incorporate guanine opposite 2,2,4-triamino-5(2H)-oxazolone. ChemBioChem, 10, 2613–2616.

95. Hunter, S.E., Gustafson, M.A., Margillo, K.M., Lee, S.A., Ryde, I.T. and Meyer, J.N. (2012) In vivo repair of alkylating and oxidative DNA damage in the mitochondrial and nuclear genomes of wild-type and glycosylase-deficient Caenorhabditis elegans. DNA Repair (Amst)., 11, 857–863.

96. SenGupta, T., Palikaras, K., Esbensen, Y.Q., Konstantinidis, G., Galindo, F.J.N., Achanta, K., Kassahun, H., Stavgiannoudaki, I., Bohr, V.A., Akbari, M., et al. (2021) Base excision repair causes age-dependent accumulation of single-stranded DNA breaks that contribute to Parkinson disease pathology. Cell Rep., 36, 109668.

97. Elsakrmy, N., Zhang-Akiyama, Q.M. and Ramotar, D. (2020) The Base Excision Repair Pathway in the Nematode Caenorhabditis elegans. Front. Cell Dev. Biol., 8, 1–15.

98. Sanada, Y. and Zhang-Akiyama, Q.M. (2014) An increase of oxidised nucleotides activates DNA damage checkpoint pathway that regulates post-embryonic development in Caenorhabditis elegans. Mutagenesis, 29, 107–114.

99. Meier, B., Cooke, S.L. and Al., E. (2014) C. elegans whole genome sequencing reveals mutational signatures related to carcinogens and DNA repair deficiency. Genome Res., 1, 1624–1636.

100. Dan, X., Babbar, M., Moore, A., Wechter, N., Tian, J., Mohanty, J.G., Croteau, D.L. and Bohr, V.A. (2020) DNA damage invokes mitophagy through a pathway involving Spata18. Nucleic Acids Res., 48, 6611–6623.

101. Vartanian, V., Minko, I.G., Chawanthayatham, S., Egner, P.A., Lin, Y.C., Earley, L.F., Makar, R., Eng, J.R., Camp, M.T., Li, L., et al. (2017) NEIL1 protects against aflatoxin-induced hepatocellular carcinoma in mice. Proc. Natl. Acad. Sci. U. S. A., 114, 4207–4212.

102. Lujan, S.A., Longley, M.J., Humble, M.H., Lavender, C.A., Burkholder, A., Blakely, E.L., Alston, C.L., Gorman, G.S., Turnbull, D.M., McFarland, R., et al. (2020) Ultrasensitive deletion detection links mitochondrial DNA replication, disease, and aging. Genome Biol., 21.

103. Chen, Z., Wang, Z.H., Zhang, G., Bleck, C.K.E., Chung, D.J., Madison, G.P., Lindberg, E., Combs, C., Balaban, R.S. and Xu, H. (2020) Mitochondrial DNA segregation and replication restrict the transmission of detrimental mutation. J. Cell Biol., 219.

104. Zhao, L. (2019) Mitochondrial DNA degradation: A quality control measure for mitochondrial genome maintenance and stress response. Enzymes, 45, 311–341.

105. Wang, Y., Zhang, Y., Chen, L., Liang, Q., Yin, X.-M., Miao, L., Kang, B.-H. and Xue, D. (2016) ARTICLE Kinetics and specificity of paternal mitochondrial elimination in Caenorhabditis elegans. Nat. Commun., 7.

106. Lim, Y., Rubio-Peña, K., Sobraske, P.J., Molina, P.A., Brookes, P.S., Galy, V. and Nehrke, K. (2019) Fndc-1 contributes to paternal mitochondria elimination in C. elegans. Dev Biol, 454, 15–20.

107. Wang, S., Tang, M., Pei, B., Xiao, X., Wang, J., Hang, H. and Wu, L. (2008) Cadmium-Induced Germline Apoptosis in Caenorhabditis elegans: The Roles of HUS1, p53, and MAPK Signaling Pathways. Toxicol. Sci., 102, 345–351.

108. Feng, W., Xue, K.S., Tang, L., Williams, P.L. and Wang, J. (2017) Aflatoxin B1-Induced Developmental and DNA Damage in Caenorhabditis elegans. Toxins (Basel)., 9.

109. U.S. Environmental Protection Agency (2021) TSCA Chemical Substance Inventory. TSCA Chem. Subst. Invent.

110. Landrigan, P.J., Fuller, R., Acosta, N.J.R., Adeyi, O., Arnold, R., Basu, N. (Nil), Baldé, A.B., Bertollini, R., Bose-O’Reilly, S., Boufford, J.I., et al. (2018) The Lancet Commission on pollution and health. Lancet, 391, 462–512.

111. Langie, S.A.S., Koppen, G., Desaulniers, D., Al-Mulla, F., Al-Temaimi, R., Amedei, A., Azqueta, A., Bisson, W.H., Brown, D.G., Brunborg, G., et al. (2015) Causes of genome instability: the effect of low dose chemical exposures in modern society. Carcinogenesis, 36 Suppl 1, S61–88.

112. Yousefzadeh, M., Henpita, C., Vyas, R., Soto-Palma, C., Robbins, P. and Niedernhofer, L. (2021) Dna damage—how and why we age? Elife, 10, 1–17.

113. Stewart, J.B. and Chinnery, P.F. (2021) Extreme heterogeneity of human mitochondrial DNA from organelles to populations. Nat. Rev. Genet., 22, 106–118.

114. Yadav, N. and Chandra, D. (2013) Mitochondrial DNA mutations and breast tumorigenesis. Biochim. Biophys. Acta, 1836, 336–44.

115. Kalsbeek, A.M.F., Chan, E.K.F., Corcoran, N.M., Hovens, C.M. and Hayes, V.M. (2017) Mitochondrial genome variation and prostate cancer: a review of the mutational landscape and application to clinical management. Oncotarget, 8, 71342–71357.

116. Kabekkodu, S.P., Bhat, S., Mascarenhas, R., Mallya, S., Bhat, M., Pandey, D., Kushtagi, P., Thangaraj, K., Gopinath, P.M. and Satyamoorthy, K. (2014) Mitochondrial DNA variation analysis in cervical cancer. Mitochondrion, 16, 73–82.

117. Chan, S.S.L. (2017) Inherited mitochondrial genomic instability and chemical exposures. Toxicology, 391, 75–83.

118. Zhao, L. and Sumberaz, P. (2020) Mitochondrial DNA Damage: Prevalence, Biological Consequence, and Emerging Pathways. Chem. Res. Toxicol., 33, 2491–2502.

