## Supplementary Materials for "Resistance of mitochondrial DNA to chemical-induced germline mutations is independent of mitophagy in *C. elegans*"

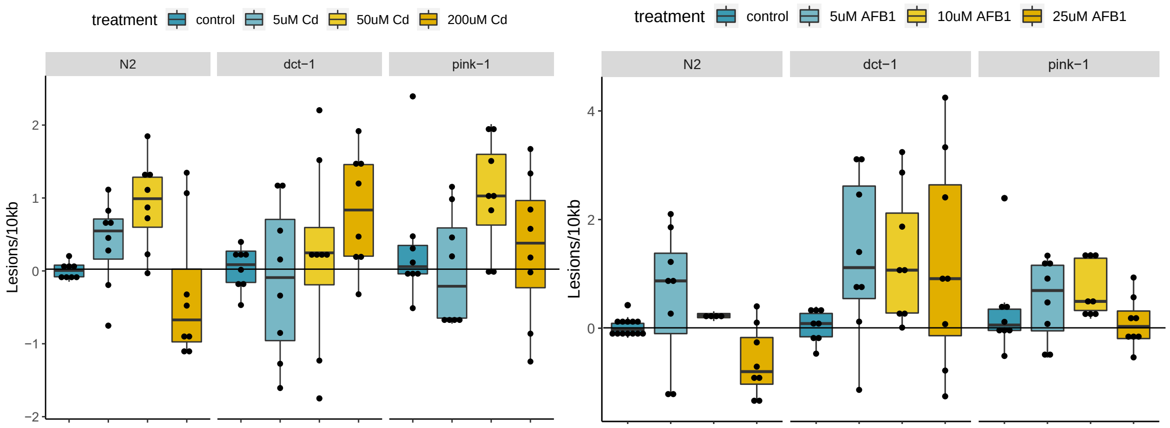


*Figure S1.* Wild-type (N2) and mitophagy mutant *C. elegans* accumulate mtDNA damage after exposure to $CdCl_2$ and $AfB_1$. We conducted a mtDNA damage dose-response for (A) $CdCl_2$ and (B) $AfB_1$ in wild-type, *dct-1*, and *pink-1* strains in order to determine a concentration for the MA line experiment that caused mtDNA lesions.

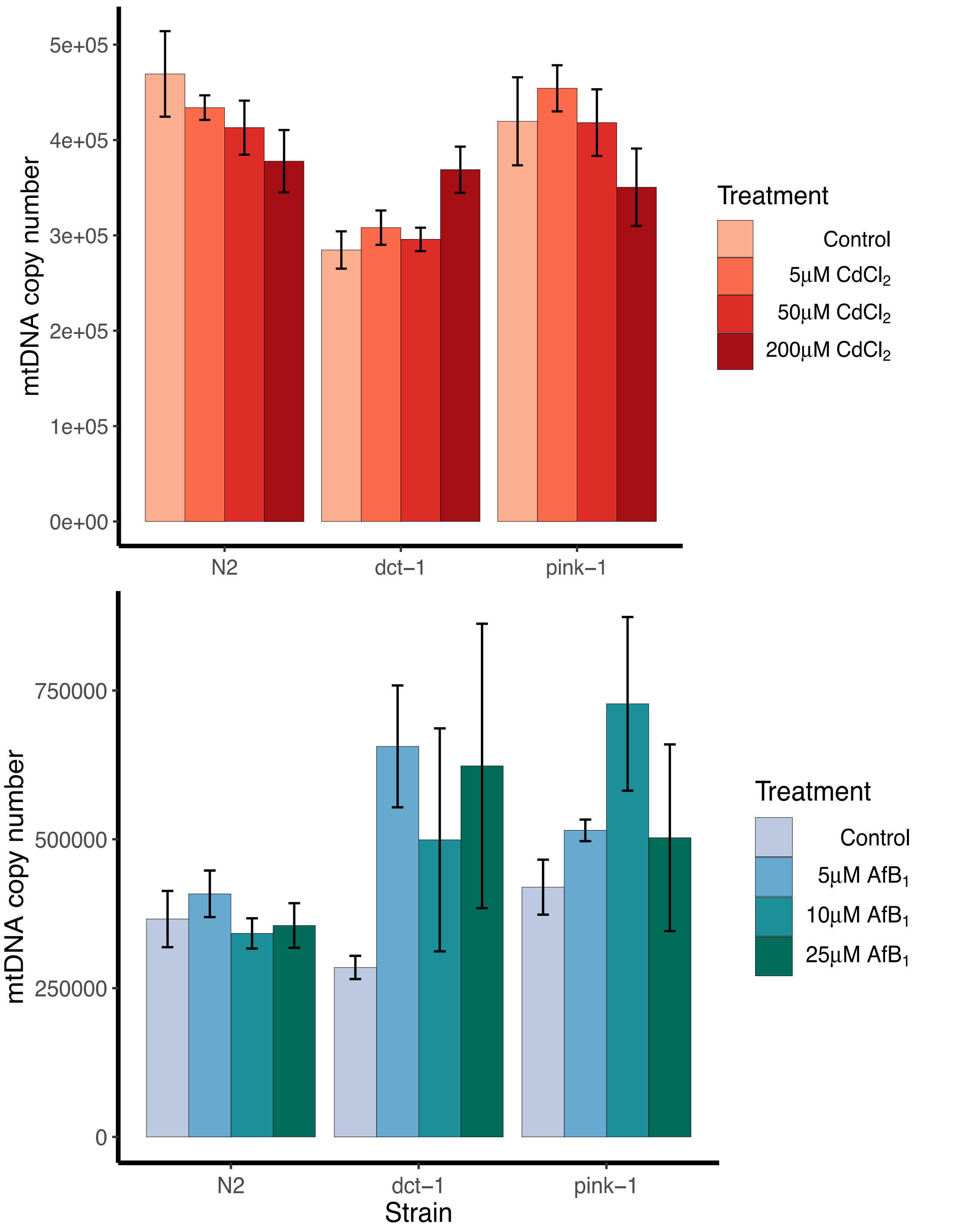


*Figure S2. mtDNA copy number quantification in L4* C. elegans *in control, $CdCl_2$, and $AfB_1$ treatments.* There is a significant strain * treatment effect on mtDNA copy number after exposure to cadmium (*P* = 0.04, two-way ANOVA, N = 8 replicates per strain per treatment). However, the mtDNA copies in the concentration that was used for the MA experiment and Duplex Sequencing (50μM) was not different from control levels in any strain. There is no significant strain * treatment effect on mtDNA copy number after exposure to aflatoxin (*P* = 0.4; two-way ANOVA; N = 4-13 replicates per strain per treatment).

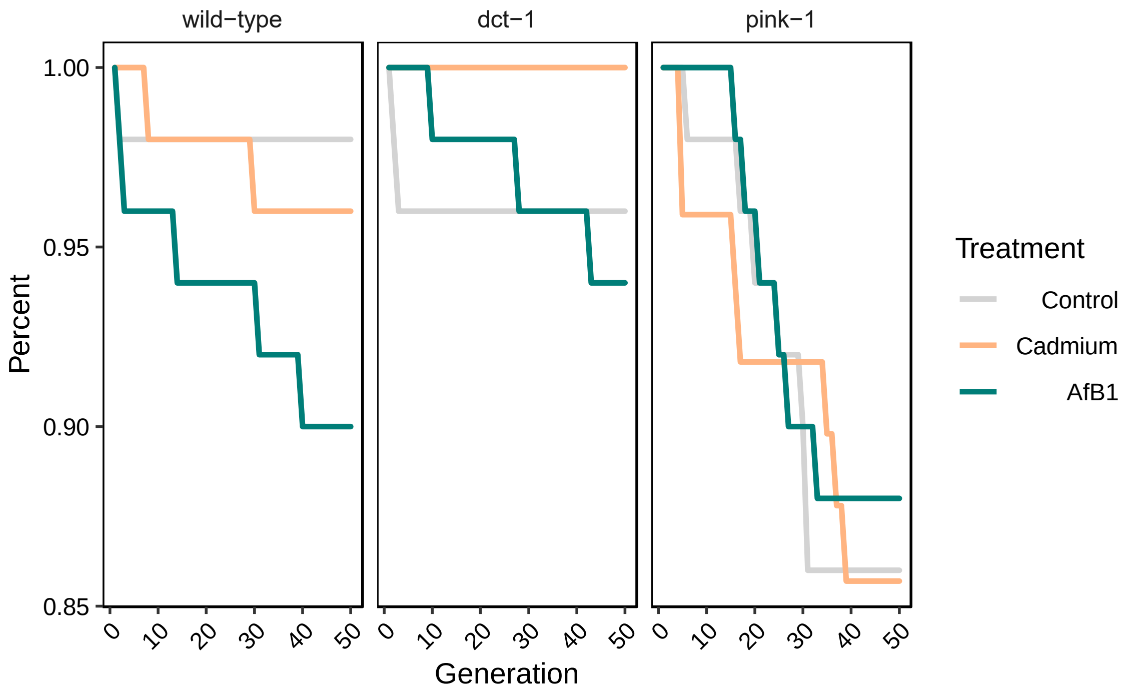


*Figure S3. Mutation accumulation line extinction rates.* The percentage of MA lines that were present at each generation out of 50 total MA lines per strain per treatment at G0. Y-axis begins at 85% survival. On average, 95.5% of MA lines were present at the end of the MA experiment (G50), though *pink-1* mutants had slightly higher extinction rates than wild-type and *dct-1* MA lines. A MA line was considered extinct if two back-up individual nematodes failed to produce offspring.

*Table S1.* Single nucleotide polymorphisms fixed in all 96 MA lines. These SNPS were not included in frequency calculations.

| Position | Ref | Alt |
|---|---|---|
| 8429 | A | G |
| 12392 | C | T |
| 12998 | C | T |

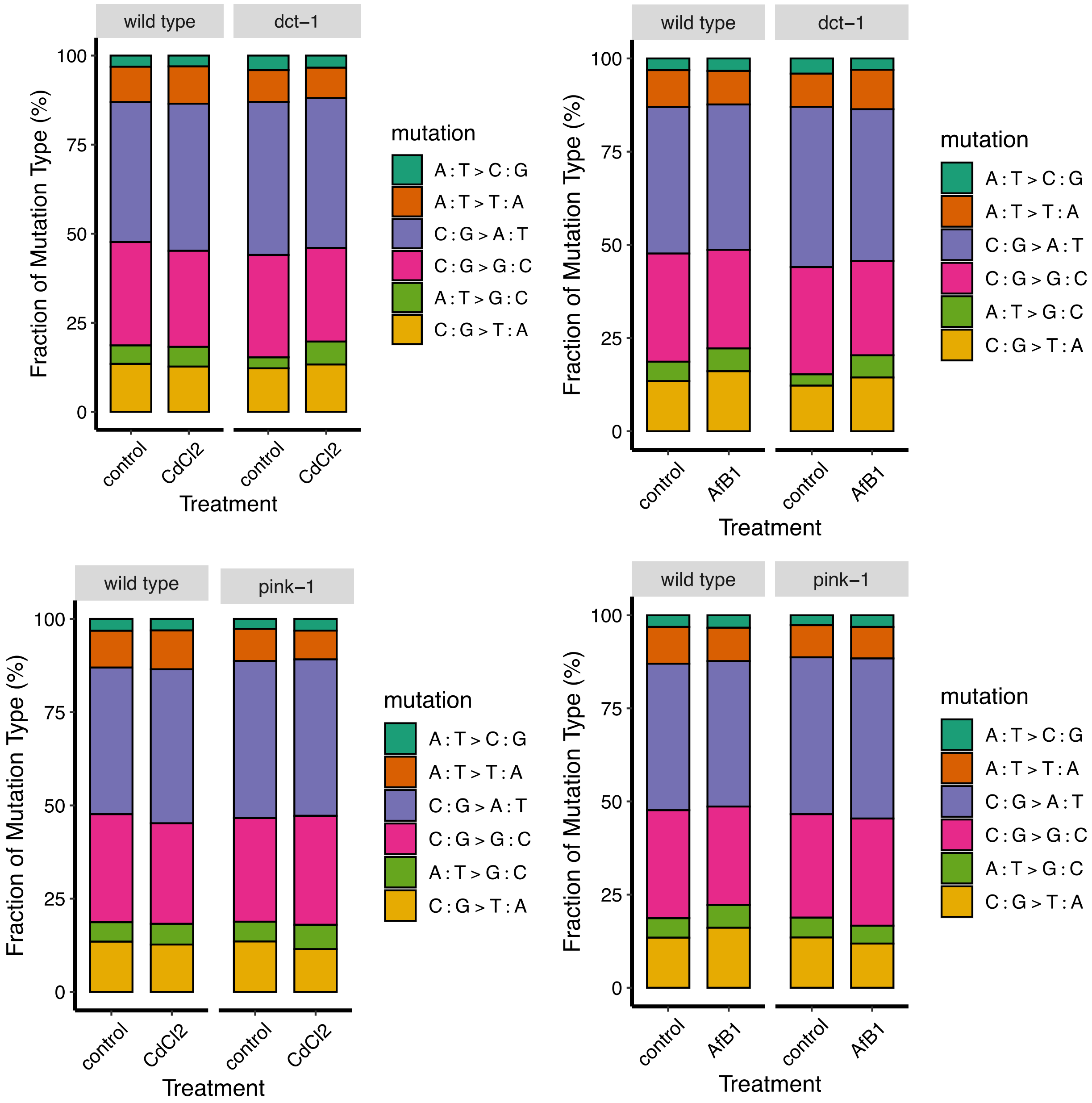


*Figure S4. Proportion of each six possible single nucleotide substitution mutations.* Composition (percentage) of each mutation type in wild-type and *dct-1* mutants in (A) cadmium and (B) aflatoxin conditions, and wild-type and *pink-1* mutants in (C) cadmium and (D) aflatoxin conditions. There is no statistical difference in the proportion of mutations between strains and no effect of treatment (Fisher's Exact Test). A majority of mutations are C:G → A:T and C:G → G:C transition mutations.

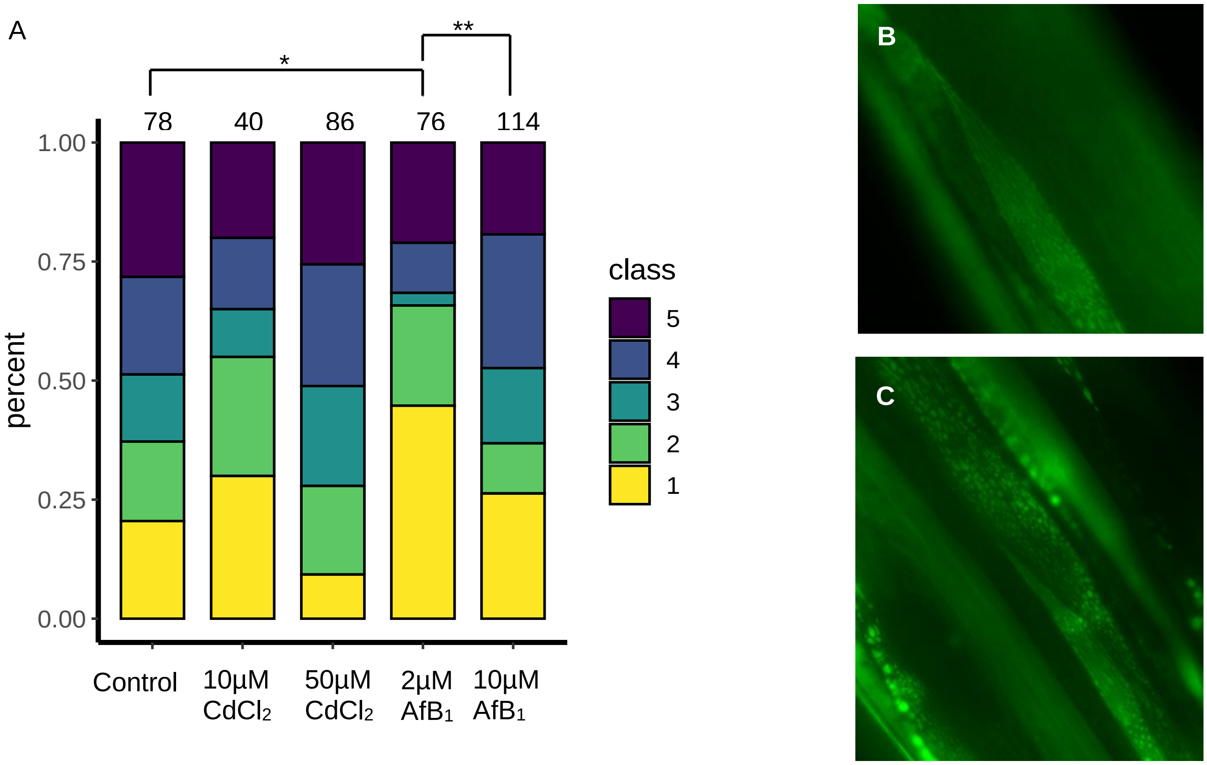


*Figure S5. Effect of cadmium and aflatoxin exposure on* C. elegans *body wall muscle mitochondrial morphology.* To assess effects of exposure on mitochondrial morphology, wild-type *C. elegans* harboring an extrachromosomal array Pmyo-3::GFP; which expresses GFP in the mitochondrial matrix of body wall muscle cells, were exposed to various concentrations of $CdCl_2$ and $AfB_1$. Images were taken on a Keyence, and analyzed in ImageJ using the Blinder Software (2). Blinded images were analyzed qualitatively by the established classification system as previously described (3), with Class 1 indicating highly networked, fused mitochondrial morphology, and Class 5 indicating extremely fragmented mitochondrial morphology. Two experimental replicates were conducted, and the total number of individuals analyzed is displayed above each stacked bar plot. (A) There were significantly fewer individuals with fragmented mitochondrial after exposure to 2µM $AfB_1$ compared to control (**$P = 0.002$; Fisher's Exact Test, Bonferroni Multiple Comparisons Correction). This suggests that mitochondria are undergoing fusion in response to a low level of stress, and fission at the higher level of $AfB_1$ exposure (***$P < 0.0002$; Fisher's Exact Test, Bonferroni Multiple Comparisons Correction). There is a trend towards a change in mitochondrial dynamics after exposure to low and high levels of $CdCl_2$ ($P = 0.02$; Fisher's Exact Test, Bonferroni Multiple Comparisons Correction), though not significant. Adjusted *alpha* = 0.005. Representative images of individuals with (B) Class 1 and (C) Class 5 mitochondrial morphology.

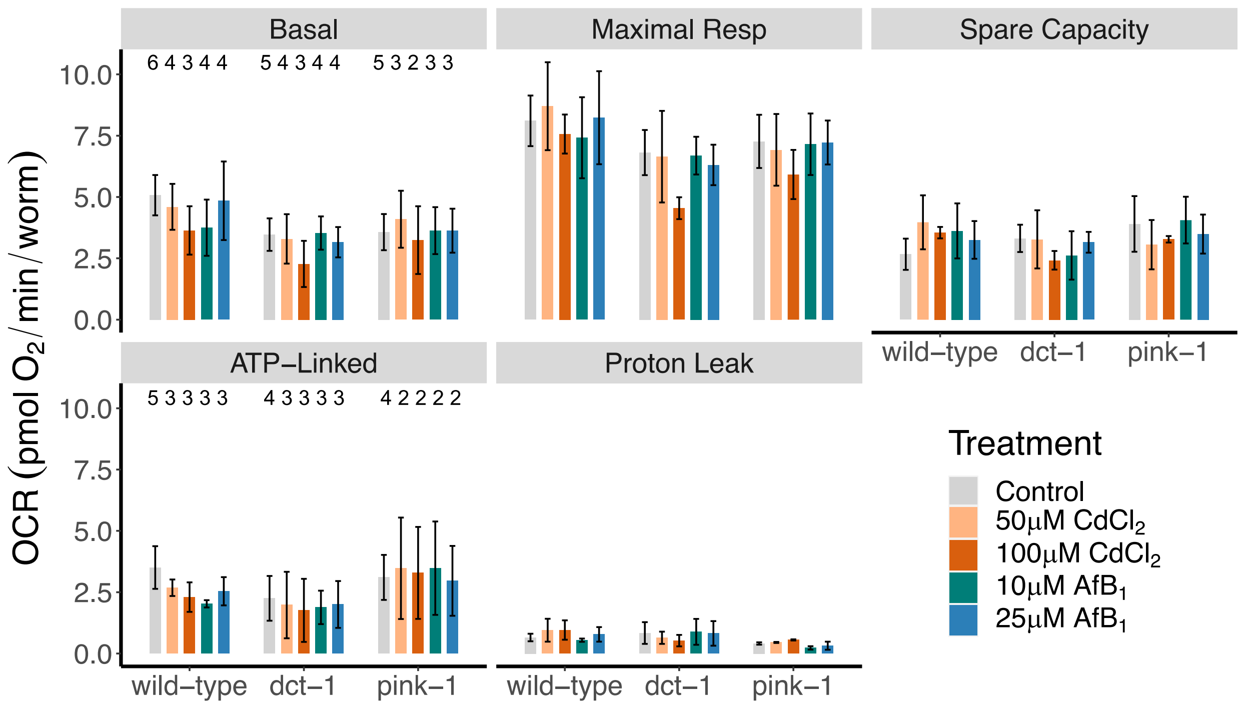


*Figure S6. Effects of cadmium and aflatoxin exposure on mitochondrial function in wild-type,* dct-1*, and* pink-1 C. elegans. Age-synchronized L1 *C. elegans* were transferred to plates seeded with OP50 containing 50µM and 100µM $CdCl_2$, or 10µM and 25µM $AfB_1$. Approximately 48 hours later, L4 *C. elegans* were washed off the plates and respiration parameters were quantified using the Seahorse Extracellular Flux Bioanalyzer, as previously described (4). The number of individual nematodes per well were counted, such that the OCR measurements were normalized per worm. The mean of technical replicates per plate was determined. Two to six independent experiments were conducted per strain per treatment. N (labeled on plot) is the same for Basal, Maximal Respiration, and Spare Capacity, and the same for ATP-Linked and Proton Leak. Two-way ANOVAs were run for each chemical in order to compare cadmium to control and aflatoxin to control across three strains. There was no significant strain * treatment interaction of cadmium compared to control in any respiration parameter ($P$ = 0.97, 0.97, 0.71, 0.97, 0.83, Basal, Maximal Respiration, and Spare Capacity, ATP-Linked, Proton Leak, respectively; two-way ANOVA) or aflatoxin compared to control in any respiration parameter ($P$ = 0.91, 0.99, 0.87, 0.90, 0.99, Basal, Maximal Respiration, and Spare Capacity, ATP-Linked, Proton Leak, respectively; two-way ANOVA).

**A**

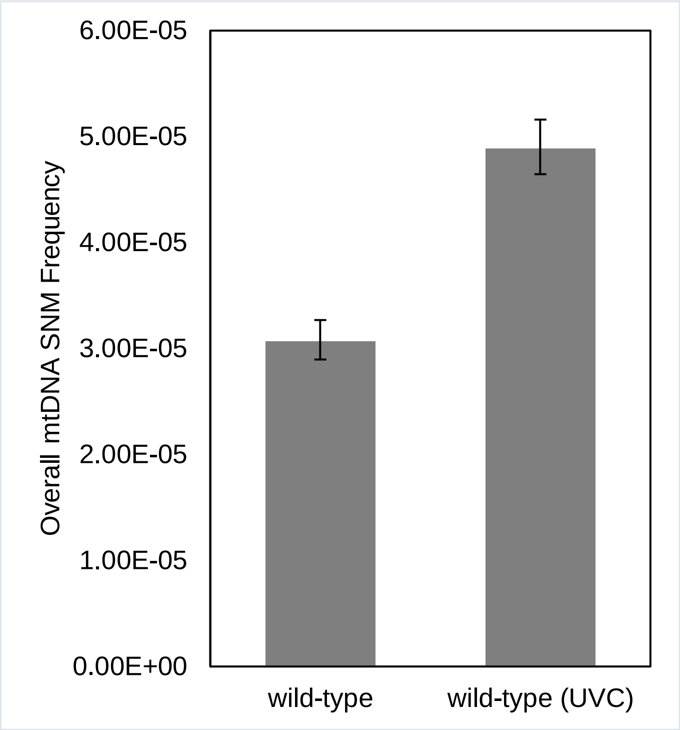


**B**

| | Control | UVC |
|---|---|---|
| Total nucleotides sequenced | 8,953,187 | 7,403,476 |
| Total point mutations: | 275 | 362 |
| Overall point mutation frequency: | 3.07E-05 | 4.89E-05 |
| 95% positive CI | 3.46E-05 | 5.42E-05 |
| 95% negative CI | 2.73E-05 | 4.41E-05 |

*Figure S7. UVC exposure results in 1.6-fold increase in mtDNA mutation frequencies in wild-type* C. elegans*.* To optimize the Duplex Sequencing library preparation and sequencing protocol for *C. elegans* mtDNA-targeted sequencing, we first sequenced a wild-type control sample and a sample of wild-type *C. elegans* that were exposure to 7 J/m$^2$ UVC radiation as a positive control. We detected hundreds of mtDNA SNMs in each sample. We observed an increase in overall mtDNA mutation frequencies. Error bars represent standard error of the mean derived from the 95% CI.

Table S2. Overall mtDNA SNM frequencies. Duplex sequencing was conducted on 9-14 MA lines per strain per treatment. There was no significant Strain x Treatment interaction in the frequency of mtDNA SNMs after exposure to 50µM $CdCl_2$ or 10µM $AfB_1$ in wild-type compared to mitophagy mutants *dct-1* and *pink-1* (two-way ANOVA, $P = 0.3$).

| Strain | Treatment | N | Mean overall SNM Frequency ($10^{-7}$) |
|---|---|---|---|
| wild-type | Control | 11 | 10.14 |
| wild-type | 50µM $CdCl_2$ | 14 | 9.75 |
| wild-type | 10µM $AfB_1$ | 10 | 9.47 |
| *dct-1* | Control | 9 | 8.93 |
| *dct-1* | 50µM $CdCl_2$ | 11 | 9.74 |
| *dct-1* | 10µM $AfB_1$ | 11 | 8.21 |
| *pink-1* | Control | 9 | 9.09 |
| *pink-1* | 50µM $CdCl_2$ | 11 | 9.16 |
| *pink-1* | 10µM $AfB_1$ | 10 | 11.63 |

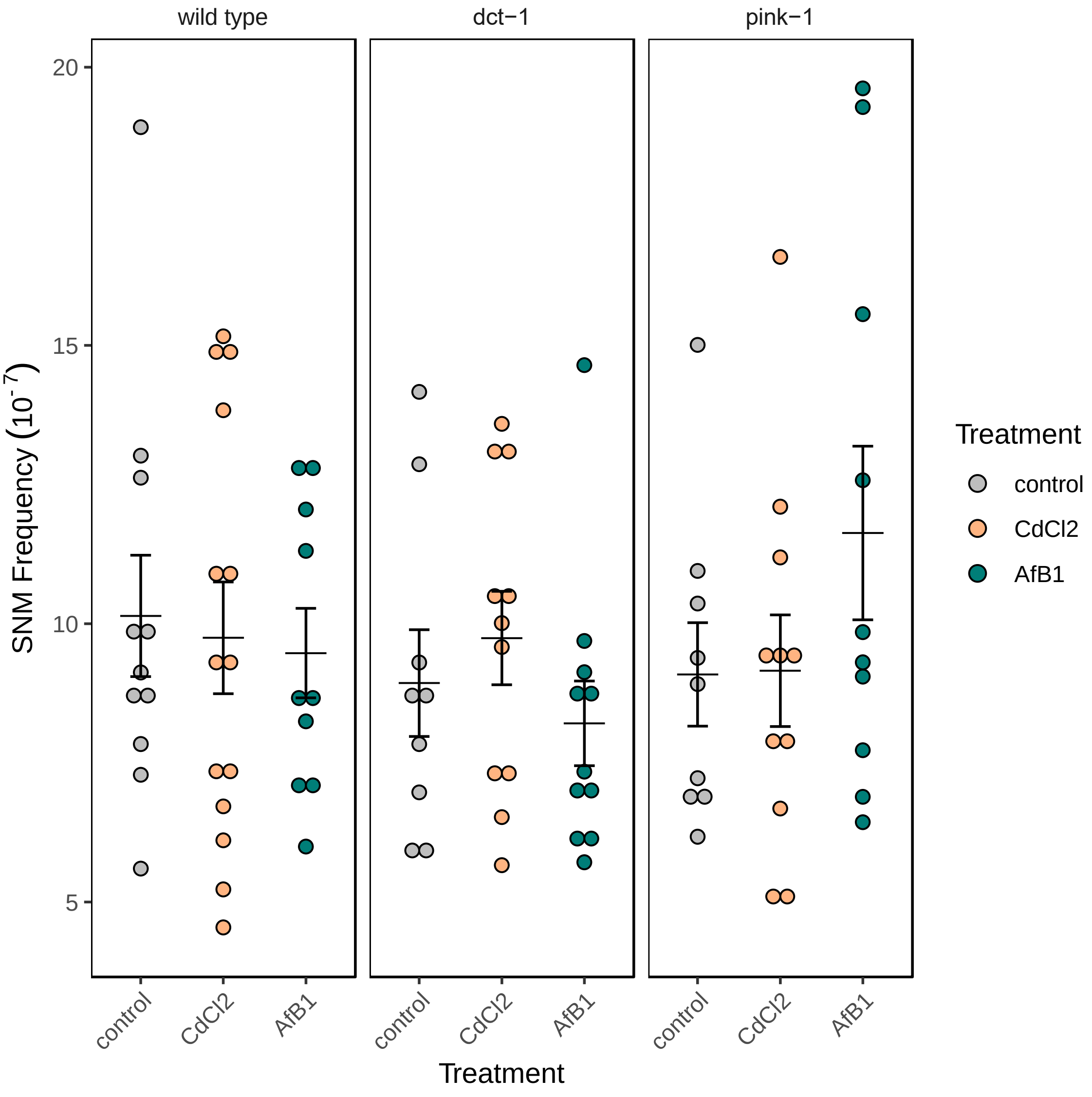


*Figure S8. Overall mtDNA SNM frequencies.* Overall SNM frequencies were determined by Duplex Sequencing after 50 generations of mutation accumulation in wild-type and two mitophagy-deficient mutants, *dct-1* and *pink-1*, *C. elegans* in control conditions (gray dots), 50µM $CdCl_2$ (gold dots), and 10µM $AfB_1$ (green dots). There was no significant effect of either 50µM $CdCl_2$ or 10µM $AfB_1$ on overall mtDNA mutation frequency compared to control between strains (two-way ANOVA, $P = 0.3$). Horizontal lines indicate mean values and error bars represent standard error of the mean.

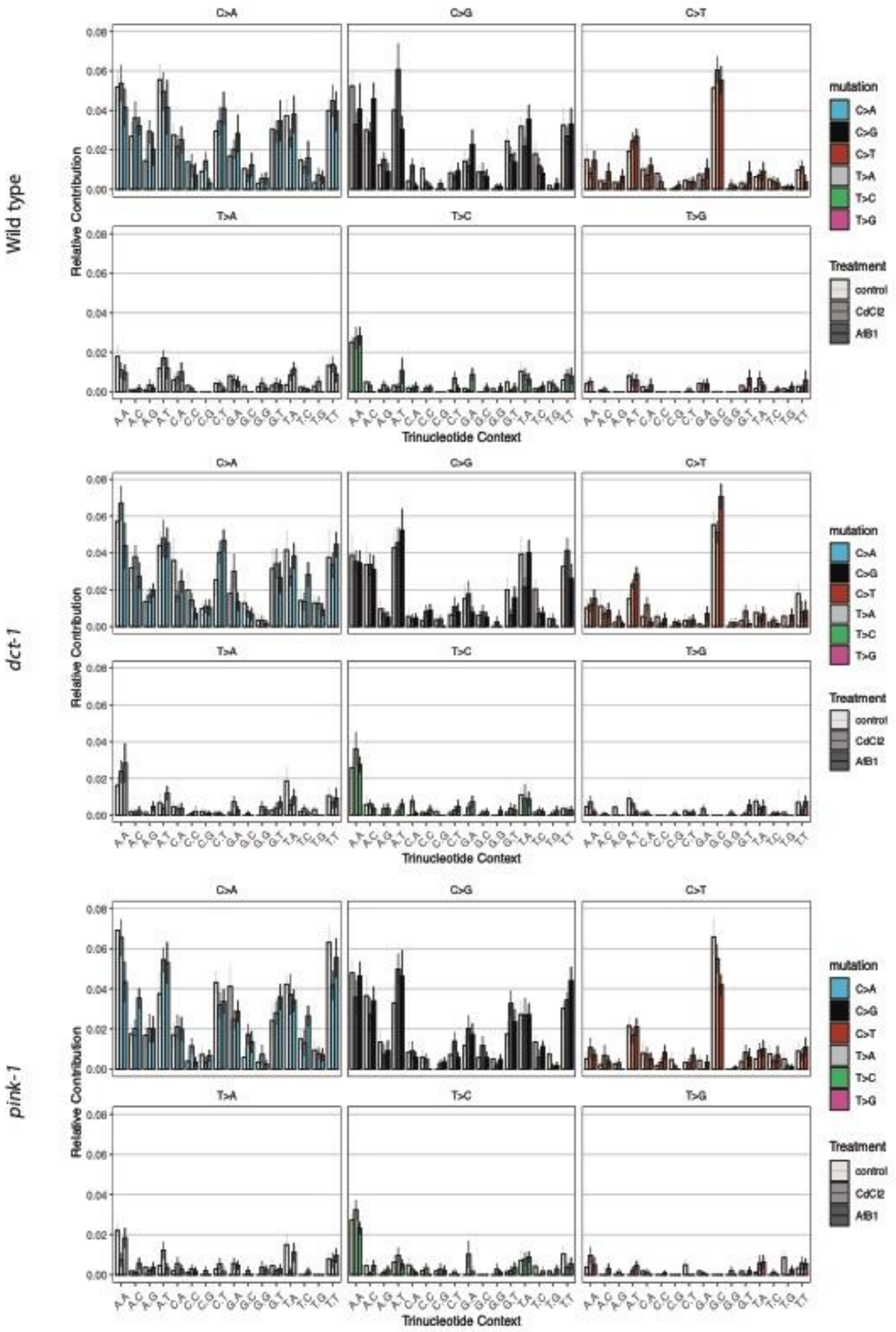


*Figure S9.* C. elegans *mtDNA trinucleotide context mutational signatures in wild-type,* dct-1*, and* pink-1*s in control conditions and $CdCl_2$ and $AfB_1$ treatment.* Contributions of each of the 96 possible trinucleotide mutations were determined for each MA line. Bar graphs show the mean relative contribution and standard error of each trinucleotide context mutation, with each mutation represented by a different color and control, $CdCl_2$, and $AfB_1$ treatment as increasing color hues.

*Table S3. Cosine similarity values of 96 single nucleotide mutational signature across MA lines*. Total mutations within strain*treatment were combined into one VCF file and the R software *MutationalPatterns* was used to create a cosine similarity matrix between all nine MA lines. All signatures shared high similarity with cosine values greater than 0.9.

| | | Wild-type | | | *dct-1* | | | *pink-1* | | |
|---|---|---|---|---|---|---|---|---|---|---|
| | | Control | CdCl2 | AfB1 | Control | CdCl2 | AfB1 | Control | CdCl2 | AfB1 |
| Wild-type | Control | 1.000 | 0.952 | 0.946 | 0.960 | 0.933 | 0.958 | 0.947 | 0.950 | 0.962 |
| | CdCl2 | 0.952 | 1.000 | 0.906 | 0.927 | 0.934 | 0.934 | 0.941 | 0.954 | 0.955 |
| | AfB1 | 0.946 | 0.906 | 1.000 | 0.953 | 0.940 | 0.946 | 0.938 | 0.938 | 0.947 |
| *dct-1* | Control | 0.960 | 0.927 | 0.953 | 1.000 | 0.931 | 0.964 | 0.936 | 0.935 | 0.957 |
| | CdCl2 | 0.933 | 0.934 | 0.940 | 0.931 | 1.000 | 0.936 | 0.926 | 0.950 | 0.949 |
| | AfB1 | 0.958 | 0.934 | 0.946 | 0.964 | 0.936 | 1.000 | 0.941 | 0.936 | 0.963 |
| *pink-1* | Control | 0.947 | 0.941 | 0.938 | 0.936 | 0.926 | 0.941 | 1.000 | 0.942 | 0.946 |
| | CdCl2 | 0.950 | 0.954 | 0.938 | 0.935 | 0.950 | 0.936 | 0.942 | 1.000 | 0.955 |
| | AfB1 | 0.962 | 0.955 | 0.947 | 0.957 | 0.949 | 0.963 | 0.946 | 0.955 | 1.000 |

*Table S4. Cosine similarity values of MA lines to COSMIC Single Base Substitution Signatures.* Using the R software package *Mutational Patterns* we investigated the similarity of MA lines to known mutational signatures. Cells in the table are colored by cosine similarity value, with those more similar to a SBS signature in red, and those less similar in blue. Overall, there is variation in similarity compared to known signatures, though these do not vary between strain and treatment.

| | Wild-type | | | *dct-1* | | | *pink-1* | | | |
|---|---|---|---|---|---|---|---|---|---|---|
| SBS Signature | Control | CdCl2 | AfB1 | Control | CdCl2 | AfB1 | Control | CdCl2 | AfB1 | Mean |
| SBS1 | 0.032 | 0.025 | 0.066 | 0.033 | 0.039 | 0.035 | 0.043 | 0.038 | 0.018 | 0.034 |
| SBS2 | 0.064 | 0.087 | 0.076 | 0.115 | 0.054 | 0.084 | 0.063 | 0.079 | 0.118 | 0.085 |
| SBS3 | 0.661 | 0.651 | 0.678 | 0.639 | 0.648 | 0.655 | 0.610 | 0.658 | 0.655 | 0.644 |
| SBS4 | 0.606 | 0.605 | 0.603 | 0.633 | 0.618 | 0.624 | 0.600 | 0.594 | 0.599 | 0.611 |
| SBS5 | 0.446 | 0.450 | 0.516 | 0.434 | 0.463 | 0.470 | 0.451 | 0.465 | 0.465 | 0.458 |
| SBS6 | 0.074 | 0.074 | 0.127 | 0.074 | 0.086 | 0.088 | 0.086 | 0.079 | 0.054 | 0.078 |
| SBS7a | 0.070 | 0.082 | 0.090 | 0.082 | 0.063 | 0.081 | 0.065 | 0.087 | 0.121 | 0.083 |
| SBS7b | 0.111 | 0.103 | 0.107 | 0.100 | 0.072 | 0.096 | 0.082 | 0.093 | 0.124 | 0.094 |
| SBS7c | 0.194 | 0.228 | 0.194 | 0.186 | 0.146 | 0.187 | 0.172 | 0.153 | 0.173 | 0.170 |
| SBS7d | 0.129 | 0.114 | 0.137 | 0.115 | 0.109 | 0.115 | 0.108 | 0.109 | 0.116 | 0.112 |
| SBS8 | 0.648 | 0.661 | 0.653 | 0.657 | 0.648 | 0.650 | 0.627 | 0.644 | 0.642 | 0.645 |
| SBS9 | 0.307 | 0.338 | 0.353 | 0.331 | 0.290 | 0.347 | 0.336 | 0.315 | 0.317 | 0.323 |
| SBS10a | 0.345 | 0.420 | 0.362 | 0.356 | 0.315 | 0.381 | 0.467 | 0.327 | 0.393 | 0.373 |
| SBS10b | 0.092 | 0.109 | 0.110 | 0.143 | 0.086 | 0.132 | 0.132 | 0.100 | 0.120 | 0.119 |
| SBS11 | 0.099 | 0.100 | 0.138 | 0.120 | 0.105 | 0.132 | 0.089 | 0.100 | 0.117 | 0.111 |
| SBS12 | 0.157 | 0.169 | 0.202 | 0.136 | 0.175 | 0.166 | 0.172 | 0.176 | 0.149 | 0.162 |
| SBS13 | 0.385 | 0.326 | 0.445 | 0.431 | 0.424 | 0.427 | 0.345 | 0.377 | 0.428 | 0.405 |
| SBS14 | 0.433 | 0.457 | 0.445 | 0.414 | 0.488 | 0.463 | 0.457 | 0.403 | 0.442 | 0.444 |
| SBS15 | 0.109 | 0.109 | 0.146 | 0.126 | 0.116 | 0.127 | 0.117 | 0.115 | 0.094 | 0.116 |
| SBS16 | 0.235 | 0.233 | 0.280 | 0.218 | 0.276 | 0.236 | 0.245 | 0.280 | 0.255 | 0.252 |
| SBS17a | 0.020 | 0.062 | 0.038 | 0.040 | 0.044 | 0.043 | 0.027 | 0.027 | 0.018 | 0.033 |
| SBS17b | 0.019 | 0.031 | 0.028 | 0.034 | 0.032 | 0.042 | 0.041 | 0.015 | 0.019 | 0.030 |
| SBS18 | 0.610 | 0.618 | 0.655 | 0.650 | 0.631 | 0.644 | 0.719 | 0.626 | 0.632 | 0.651 |
| SBS19 | 0.112 | 0.112 | 0.149 | 0.102 | 0.094 | 0.105 | 0.095 | 0.099 | 0.133 | 0.105 |
| SBS20 | 0.253 | 0.262 | 0.277 | 0.234 | 0.305 | 0.277 | 0.243 | 0.226 | 0.212 | 0.250 |
| SBS21 | 0.082 | 0.078 | 0.136 | 0.065 | 0.103 | 0.107 | 0.083 | 0.111 | 0.071 | 0.090 |
| SBS22 | 0.137 | 0.156 | 0.142 | 0.149 | 0.120 | 0.153 | 0.121 | 0.120 | 0.126 | 0.132 |
| SBS23 | 0.092 | 0.093 | 0.127 | 0.090 | 0.084 | 0.110 | 0.090 | 0.091 | 0.094 | 0.093 |
| SBS24 | 0.439 | 0.427 | 0.457 | 0.465 | 0.472 | 0.462 | 0.477 | 0.459 | 0.466 | 0.467 |
| SBS25 | 0.421 | 0.441 | 0.472 | 0.426 | 0.428 | 0.453 | 0.417 | 0.416 | 0.423 | 0.427 |
| SBS26 | 0.117 | 0.129 | 0.165 | 0.098 | 0.151 | 0.132 | 0.133 | 0.146 | 0.113 | 0.129 |
| SBS28 | 0.055 | 0.056 | 0.076 | 0.069 | 0.056 | 0.086 | 0.053 | 0.060 | 0.064 | 0.065 |
| SBS29 | 0.569 | 0.563 | 0.600 | 0.611 | 0.635 | 0.590 | 0.625 | 0.603 | 0.594 | 0.610 |
| SBS30 | 0.135 | 0.129 | 0.193 | 0.124 | 0.131 | 0.149 | 0.108 | 0.130 | 0.150 | 0.132 |
| SBS31 | 0.223 | 0.217 | 0.227 | 0.197 | 0.191 | 0.215 | 0.201 | 0.205 | 0.232 | 0.207 |
| SBS32 | 0.212 | 0.212 | 0.302 | 0.225 | 0.254 | 0.259 | 0.204 | 0.220 | 0.236 | 0.233 |
| SBS33 | 0.104 | 0.106 | 0.097 | 0.075 | 0.079 | 0.091 | 0.086 | 0.079 | 0.102 | 0.085 |
| SBS34 | 0.169 | 0.188 | 0.186 | 0.183 | 0.113 | 0.180 | 0.158 | 0.119 | 0.146 | 0.150 |
| SBS35 | 0.402 | 0.404 | 0.411 | 0.402 | 0.412 | 0.407 | 0.364 | 0.402 | 0.415 | 0.400 |
| SBS36 | 0.571 | 0.557 | 0.608 | 0.600 | 0.557 | 0.618 | 0.670 | 0.549 | 0.592 | 0.598 |

| | | | | | | | | | | |
|---|---|---|---|---|---|---|---|---|---|---|
| SBS37 | 0.280 | 0.277 | 0.310 | 0.254 | 0.283 | 0.274 | 0.292 | 0.286 | 0.265 | 0.276 |
| SBS38 | 0.283 | 0.292 | 0.292 | 0.345 | 0.288 | 0.308 | 0.248 | 0.269 | 0.265 | 0.287 |
| SBS39 | 0.679 | 0.640 | 0.666 | 0.643 | 0.641 | 0.652 | 0.619 | 0.671 | 0.671 | 0.650 |
| SBS40 | 0.694 | 0.695 | 0.734 | 0.696 | 0.694 | 0.707 | 0.681 | 0.696 | 0.711 | 0.697 |
| SBS41 | 0.317 | 0.344 | 0.366 | 0.344 | 0.282 | 0.337 | 0.314 | 0.297 | 0.324 | 0.316 |
| SBS42 | 0.263 | 0.269 | 0.303 | 0.276 | 0.275 | 0.309 | 0.274 | 0.260 | 0.271 | 0.277 |
| SBS44 | 0.252 | 0.256 | 0.314 | 0.218 | 0.275 | 0.272 | 0.244 | 0.229 | 0.209 | 0.241 |
| SBS84 | 0.215 | 0.208 | 0.265 | 0.207 | 0.219 | 0.226 | 0.201 | 0.244 | 0.220 | 0.219 |
| SBS85 | 0.306 | 0.313 | 0.305 | 0.289 | 0.259 | 0.320 | 0.293 | 0.244 | 0.273 | 0.280 |
| SBS86 | 0.542 | 0.526 | 0.537 | 0.531 | 0.544 | 0.523 | 0.493 | 0.560 | 0.547 | 0.533 |
| SBS87 | 0.156 | 0.156 | 0.174 | 0.164 | 0.166 | 0.152 | 0.176 | 0.163 | 0.152 | 0.162 |
| SBS88 | 0.189 | 0.198 | 0.247 | 0.170 | 0.182 | 0.217 | 0.212 | 0.210 | 0.198 | 0.198 |
| SBS89 | 0.624 | 0.618 | 0.668 | 0.617 | 0.637 | 0.640 | 0.591 | 0.629 | 0.623 | 0.623 |
| SBS90 | 0.127 | 0.173 | 0.169 | 0.186 | 0.115 | 0.168 | 0.157 | 0.091 | 0.155 | 0.145 |

*Table S5. Cosine similarity values to COSMIC Single Base Substitution Signatures.* The mean cosine similarity value across all nine MA lines was determined for each of the 54 SBS Signatures from the COSMIC database (Table S4). Here the SBS signatures are ranked from high (similar) to low (not similar) cosine similarity values. There are 10 mutational signatures that have a cosine similarity value >0.5. The proposed etiology, if known, is listed.

| SBS Signature | Mean Cosine Similarity Value | Proposed Etiology |
|---|---|---|
| SBS40 | 0.701 | unknown (age) |
| SBS39 | 0.654 | unknown |
| SBS3 | 0.651 | Defective homologous recombination-based DNA damage repair, abnormal double strand break repair |
| SBS8 | 0.648 | unknown |
| SBS18 | 0.643 | **ROS** |
| SBS89 | 0.627 | unknown |
| SBS4 | 0.609 | Tobacco chewing |
| SBS29 | 0.599 | Tobacco smoking |
| SBS36 | 0.591 | Defective base excision repair, including DNA damage due to reactive oxygen species, due to biallelic germline or somatic *MUTYH* mutations. |
| SBS86 | 0.534 | Unknown chemotherapy treatment |
| SBS5 | 0.462 | Unknown (clock-like signature) |
| SBS24 | 0.458 | **Aflatoxin exposure** |
| SBS14 | 0.445 | Pol Epsilon mutation and defective MMR |
| SBS25 | 0.433 | Chemotherapy |
| SBS35 | 0.402 | Platinum chemotherapy |
| SBS13 | 0.399 | APOBEC deaminases |
| SBS10a | 0.374 | Pol Epsilon exonuclease domain mutations |
| SBS9 | 0.326 | Pol eta hypermutation |
| SBS41 | 0.325 | unknown |
| SBS85 | 0.289 | Indirect effects of activation-induced cytidine deamination |
| SBS38 | 0.288 | UV (indirect) |
| SBS37 | 0.280 | unknown |
| SBS42 | 0.278 | Haloalkane exposure |
| SBS20 | 0.254 | Defective MMR and POLD1 mutations |
| SBS44 | 0.252 | Defective MMR |
| SBS16 | 0.251 | unknown |
| SBS32 | 0.236 | Azathioprine treatment |
| SBS84 | 0.223 | Activation-induced cytidine deamination |
| SBS31 | 0.212 | Platinum chemotherapy |
| SBS88 | 0.203 | Colibactin exposure |
| SBS7c | 0.182 | UV exposure |
| SBS12 | 0.167 | unknown |
| SBS87 | 0.162 | Thiopurine chemotherapy |
| SBS34 | 0.160 | unknown |
| SBS90 | 0.149 | Duocarmycin exposure |
| SBS30 | 0.139 | Defective BER (NTHL-1 mutations) |
| SBS22 | 0.136 | Aristolochic Acid exposure |
| SBS26 | 0.132 | Defective MMR |
| SBS15 | 0.118 | Defective MMR |

| | | |
|---|---|---|
| SBS7d | 0.117 | UV exposure |
| SBS10b | 0.114 | Pol Epsilon exonuclease domain mutations |
| SBS19 | 0.111 | unknown |
| SBS11 | 0.111 | Temozolomide treatment |
| SBS7b | 0.099 | UV exposure |
| SBS23 | 0.097 | unknown |
| SBS21 | 0.093 | Defective MMR |
| SBS33 | 0.091 | unknown |
| SBS6 | 0.083 | Defective MMR |
| SBS7a | 0.082 | UV exposure |
| SBS2 | 0.082 | APOBEC deaminases |
| SBS28 | 0.064 | unknown |
| SBS1 | 0.037 | 5-methylcytosine deamination |
| SBS17a | 0.035 | unknown |
| SBS17b | 0.029 | unknown |

*Table S6. Mutation accumulation lines are under neutral evolution.* R package *dNdScv* was used to calculate the dN/dS ratio across wild-type, *dct-1*, and *pink-1* MA lines across control, cadmium, and aflatoxin treatment. Wmis = missense mutations, wnon = nonsense mutations, wall = all nonsynonymous mutations. All predicted global dN/dS values across all strains/treatments are close to 1, suggesting that the *C. elegans* mitochondrial genome is under neutral evolution in the MA experimental design.

| Strain | Treatment | Name | Maximum likelihood model | CI (low) | CI (high) |
|---|---|---|---|---|---|
| wildtype | control | wmis | 0.924 | 0.702 | 1.216 |
| | | wnon | 0.908 | 0.542 | 1.520 |
| | | wall | 0.924 | 0.702 | 1.216 |
| wildtype | cadmium | wmis | 1.159 | 0.887 | 1.515 |
| | | wnon | 0.958 | 0.579 | 1.585 |
| | | wall | 1.155 | 0.884 | 1.509 |
| wildtype | aflatoxin | wmis | 1.1231009 | 0.8322949 | 1.515515 |
| | | wnon | 0.7382623 | 0.3984926 | 1.367732 |
| | | wall | 1.1151789 | 0.8263014 | 1.505049 |
| *dct-1* | control | wmis | 0.956759 | 0.6895992 | 1.32742 |
| | | wnon | 0.6145301 | 0.3173671 | 1.189938 |
| | | wall | 0.9465993 | 0.6821831 | 1.313504 |
| *dct-1* | cadmium | wmis | 0.9454341 | 0.7281853 | 1.227498 |
| | | wnon | 0.7091876 | 0.3868122 | 1.300236 |
| | | wall | 0.942165 | 0.7256459 | 1.223289 |
| *dct-1* | aflatoxin | wmis | 1.0212917 | 0.7568162 | 1.37819 |
| | | wnon | 0.8483719 | 0.4574501 | 1.573362 |
| | | wall | 1.0180673 | 0.7544422 | 1.373811 |
| *pink-1* | control | wmis | 0.812797 | 0.600292 | 1.100529 |
| | | wnon | 0.5416891 | 0.2885511 | 1.016898 |
| | | wall | 0.8070964 | 0.5959419 | 1.093067 |
| *pink-1* | cadmium | wmis | 1.209978 | 0.8946023 | 1.636534 |
| | | wnon | 1.209296 | 0.6848774 | 2.135269 |
| | | wall | 1.209968 | 0.8946646 | 1.636394 |
| *pink-1* | aflatoxin | wmis | 1.0284231 | 0.7750509 | 1.364625 |
| | | wnon | 0.6751625 | 0.3787332 | 1.203603 |
| | | wall | 1.0195962 | 0.7683399 | 1.353016 |

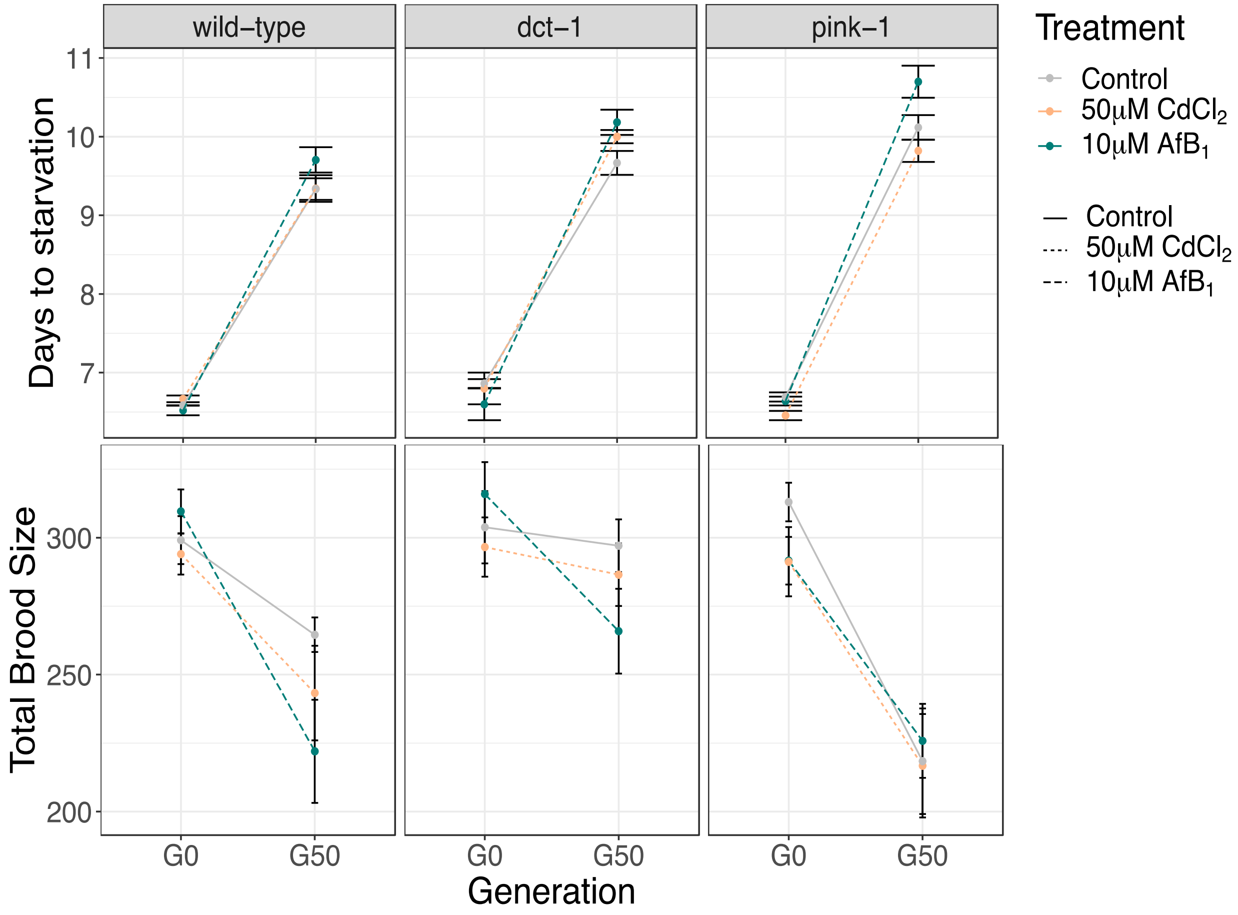


*Figure S10. Mutation accumulation results in fitness decline.* (A) We measured population growth rate as an indicator of fitness in ancestors (G0) and after 50 generations of MA in all lines. Each dot represents the mean value of days to starvation, with error bars as standard error of the mean (N = 3 for G0, N = 39-48 for G50 MA lines). Lines indicate the rate of change in fitness from G0 to G50 (gray solid = control, gold small dash = 50µM $CdCl_2$, and green large dash = 10µM $AfB_1$) for wild-type, *dct-1*, and *pink-1* strains. Y-axis lower limit begins at Day 6. Plates were monitored hourly for starvation, or until all of the food was eliminated and the population dispersed on the plate. There was no effect of $CdCl_2$ or 10µM $AfB_1$ on population growth rate in any strain at G0. All MA lines had significantly slower population growth rate at G50 compared to G0 (two-way ANOVA, $P < 2.2e\text{-}16$). There was no effect of either $CdCl_2$ or 10µM $AfB_1$ on wild-type MA lines at G50 ($P$ = 1, $P$ = 0.95, respectively; two-way ANOVA, Tukey HSD). There was no effect of either $CdCl_2$ or 10µM $AfB_1$ on *dct-1* MA lines at G50 ($P$ = 0.98, $P$ = 0.6, respectively; two-way ANOVA, Tukey HSD). There was no effect of either $CdCl_2$ or 10µM $AfB_1$ on *pink-1* MA lines at G50 ($P$ = 1, $P$ = 0.4, respectively; two-way ANOVA, Tukey HSD). Both control and 10µM $AfB_1$ *pink-1* MA lines had a significantly greater decline in fitness compared to control and 10µM $AfB_1$ wild-type MA lines ($P$ = 0.03, $P$ = .0008, respectively; two-way ANOVA, Tukey HSD). This decline in fitness suggests that *pink-1* MA lines are accumulating more deleterious mutations than wild-type MA lines. (B) Total brood size was quantified in 20 randomly selected MA lines per strain per treatment. At G50, three individual L4s per MA line were transferred onto an individual control plate. Each individual was transferred every day until cessation of egg-laying. Reproduction was counted on the previous plate after 48 hours. Total reproduction was calculated per plate. The mean of each biological replicate per MA line was then calculated. Each dot represents the mean value total brood size per MA line, with error bars indicating standard error of the mean (N = 9-10 for G0, N = 18-20 for G50 MA lines). Lines indicate the rate of change in fitness from G0 to G50 (gray solid = control, gold small dash = 50µM $CdCl_2$, and green large dash = 10µM $AfB_1$) for wild-type, *dct-1*, and *pink-1* strains. The y-axis lower limit begins at 190. All MA lines had significantly lower fecundity at G50 compared to G0 ($P$ = 2.596e-11; two-way ANOVA). However, there was no effect of either $CdCl_2$ or 10µM $AfB_1$ on total brood size in wild-type MA lines ($P$ =

1, $P = 0.7$, respectively ; two-way ANOVA, Tukey HSD). *pink-1* control MA lines had significantly smaller broods than *dct-1* control MA lines at G50 ($P < 0.01$; two-way ANOVA, Tukey HSD) and *pink-1* $CdCl_2$ MA lines had significantly smaller broods than *dct-1* $CdCl_2$ MA lines at G50 ($P = 0.03$), suggesting a decrease in fitness in *pink-1* MA lines compared to *dct-1* MA lines.

*Table S7. Internal body burden of $CdCl_2$ in wild-type* C. elegans. We measured the body burden of *$CdCl_2$* in adult *C. elegans* after a chronic exposure to 50µM $CdCl_2$. The actual concentration of Cd in the OP50 food source was slightly lower than 50µM (28 µM). However, the geometric mean of the internal concentration detected in three biological replicates averaged to 0.54 µg/L, which is comparable to human blood Cd levels (1), and 4-fold greater than control *C. elegans* samples.

| | | | $[Cd]_{ICP}$ | $[Cd]_{worms}$ | $[Cd]_{worms}$ | $[Cd]_{expected}$ | % difference | |
|---|---|---|---|---|---|---|---|---|
| | | | µg/L | %RSD | µg/L | µM | µg/L | Mean |
| | digestion blank1 | Blank | 0 | | | | | |
| | digestion blank2 | Blank | 0 | | | | | |
| | SRM | SRM | 3.6 | 2.7 | 10.7 | | 10 | 7.3 |
| control | 500 worms | | 3.3 | 3.0 | 69.6 | 0.62 | 0.139 | 0.142 |
| control | 500 worms | | 3.3 | 3.2 | 69.0 | 0.61 | 0.138 | |
| control | 1000 worms | | 3.7 | 1.6 | 73.7 | 0.66 | 0.147 | |
| 50µM $CdCl_2$ | 500 worms | | 13.8 | 1.6 | 299.6 | 2.67 | 0.599 | 0.541 |
| 50µM $CdCl_2$ | 500 worms | | 14.0 | 1.1 | 296.8 | 2.64 | 0.594 | |
| 50µM $CdCl_2$ | 1000 worms | | 21.4 | 1.5 | 430.9 | 3.83 | 0.431 | |
| 50µM $CdCl_2$ | OP50 - spiked | | 154.1 | 0.6 | 3148.9 | 28.0 | | |
